# The molecular organization of flat and curved caveolae indicates bendable structural units at the plasma membrane

**DOI:** 10.1101/2022.03.31.486578

**Authors:** Claudia Matthaeus, Kem A. Sochacki, Andrea Dickey, Dmytro Puchkov, Volker Haucke, Martin Lehmann, Justin W. Taraska

## Abstract

Caveolae are small coated inner plasma membrane invaginations found in many cell types. Their diverse functions span from endocytosis to signaling, regulating key cellular processes including lipid uptake, pathogen entry, and membrane tension. Caveolae undergo shape changes from flat to curved. It is unclear which proteins regulate this process. To address this gap, we studied the shapes of caveolae with platinum replica electron microscopy in six common cell types. Next, we developed a correlative multi-color stimulated emission depletion (STED) fluorescence and platinum replica EM imaging (CLEM) method to image caveolae-associated proteins at caveolae of different shapes at the nanoscale. Caveolins and cavins were found at all caveolae, independent of their curvature. EHD2, a classic caveolar neck protein, was strongly detected at both curved and flat caveolae. Both pacsin2 and the regulator EHBP1 were found only at a subset of caveolae. Pacsin2 was localized primarily to areas surrounding flat caveolae, whereas EHBP1 was mostly detected at spheres. Contrary to classic models, dynamin was absent from caveolae and localized only to clathrin-coated structures. Cells lacking dynamin showed no substantial changes to caveolae, suggesting that dynamin is not directly involved in caveolae curvature. Together, we provide a mechanistic map for the molecular control of caveolae shape by eight of the major caveolae-associated coat and regulatory proteins. We propose a model where caveolins, cavins, and EHD2 assemble as a cohesive structural unit regulated by more intermittent associations with pacsin2 and EHBP1. These complexes can flatten and curve, capturing membrane to enable lipid traffic and changes to the surface area of the cell.

## Introduction

Caveolae are 60-100 nm diameter coated plasma membrane domains that invaginate into the cytosol. They are prominent and common features of the plasma membrane in many cells including adipocytes, fibroblast, and muscle cells^1^. They concentrate specific lipids including cholesterol, sphingolipids, and PI(4,5)P_2_, supporting the clustering of distinct proteins and signaling molecules^2,3^. Mice lacking caveolae have defects in lipid uptake, blood vessel function, and membrane tension regulation^4^. Dysregulation or mutation of caveolae proteins are known drivers of diseases including muscle disorders^5,6^ and cancer^7^.

Much is known about the molecular components of caveolae. The caveolin proteins (caveolin1-3 in mammals) and cavins (cavin1-4) are central for organelle formation. Deletion of either caveolin1 or cavin1 results in loss of caveolae *in vivo*^4^. Structural data indicated that cavins form hetero-trimers (mainly cavin1/2 or cavin1/3) and assemble as a layered protein coat with caveolins^8–10^. Membrane remodeling proteins such as Eps15 homology (EH) domain-containing protein 2 (EHD2)^11,12^, Pacsin/Syndapin2^13–15^ and EH domain-binding protein 1 (EHBP1^16^) also associate with caveolae. In particular, the ATPase EHD2 plays an important role in stabilizing caveolae at the plasma membrane and regulating endocytosis^11,12^. Loss of EHD2 *in vivo* did not change caveolae assembly and number. However, an increased mobility of caveolae was detected in mice lacking EHD2 that was associated with an increase in lipid accumulation^17^. EHD2 has been localized to the caveolae neck with immunogold electron microscopy of thin sections^18^. Based on structural data, EHD2 has been proposed to specifically form a ring-like oligomer encircling the caveolar neck^19^. The BAR domain containing protein pacsin2 was also found at the caveolae neck^15^. Deletion of the muscle-specific variant pacsin3 led to a loss of the characteristic caveolae bulb shape despite the fact that caveolin1 and cavin1 were still present at the plasma membrane^20^. Knockdown of pacsin2 leads to shallow caveolae, and impairs caveolae mobility and endocytosis^13,15,21^. Recently, EHBP1 was found to stabilize caveolae at the plasma membrane. Loss of EHBP1, similar to loss of EHD2, however, did not modulate caveolae shapes but rather increased endocytosis^16^. Additionally, dynamin has been commonly implicated in caveolae endocytosis and proposed to have a similar role to its well-established functions in membrane scission in clathrin-mediated endocytosis^22–24^. Other notable proteins have been linked to caveolae including receptor tyrosine kinase-like orphan receptor 1 (ROR1) in embryonic tissues^25^ or the c-Abl tyrosine kinase FBP17 in rosette-like caveolae clusters^26,27^. Yet, while much data is known, it is unclear how these components assemble and curve at the plasma membrane^28,29^. Understanding this architecture is necessary for understanding how caveolae function in cells, what their roles are, and how they are regulated across different pathways and tissues.

The structures of purified caveolin1 complexes suggest that the assembly of several caveolin1 molecules into ∼14 nm sized disc-shaped complexes is needed to induce caveolae formation^30–32^. Cavins are then recruited to these sites to leading to induce substantial membrane bending. This process is proposed to be reversible during increased membrane tension (e.g. osmotic shock) or cellular stress (e.g. UV light) where cavins are released, leading to a flattening of the invagination^10,33,34^. However, it is currently unknown how flat and deeply curved caveolae differ in their morphologies and protein components. Additionally, it is unclear if cavins are completely lost during flatting, if caveolae disassemble upon increased membrane tension, or if caveolae exhibit a more flexible coat complex that, similar to clathrin-coated pits, can change its shape from flat to curved. Furthermore, it is not known when, if, and how the caveolae proteins EHD2, pacsin2 and EHBP1 are recruited to these sites and how these proteins regulate the caveolin/cavin coat complex and its shape. As an example, EHD2 was shown to translocate into the nucleus after caveolae flatten^35^ indicating that EHD2 may not associate with flat caveolae. A detailed understanding of the coat and associated proteins, however, in relation to caveolae shape and curvature is missing. These questions have been difficult to answer with light microscopy due to the small size of caveolae relative to the diffraction limit. Also, past electron microscopy (EM) measurements were insufficient for two reasons. First, imaging all caveolae in a membrane to provide a population-level structural view is challenging in thin section EM, and second, localizing and quantifying specific components within those EM images is difficult with established labeling and analysis methods.

To overcome these limitations, we investigate the relationship between caveolae morphology and the proteins classically proposed to regulate caveolae structure and behavior with nanoscale correlative light and electron microscopy across entire plasma membranes. First, to understand the shape of caveolae across single cells, we use platinum replica electron microscopy (PREM) to classify and analyze visible caveolae membrane domains at the plasma membrane into flat, bulb, and spherical caveolae. Next, to understand how proteins associate with these domains, we develop a super-resolution STED and platinum replica correlation method (STED-CLEM) to localize major caveolae coat and regulatory proteins in and around single caveolae across entire plasma membranes of cultured mammalian cells. Surprisingly, in contrast to previous models, we find that along with caveolins, EHD2 and cavins were present on flat and curved caveolae equally, while EHBP1 was mainly found at a subset of curved caveolae. Pacsin2 was primarily detected at flat caveolae. Dynamin was absent from caveolae. Loss of these proteins differentially affected caveolae shape and abundance. Taken together, we present direct nanoscale insights into the control of caveolae shape across the plasma membrane and propose a new molecular model for the control of caveolae curvature in mammalian cells.

## Results

### Structural investigation and classification of membrane curvature in caveolae

Caveolae coats are proposed to change their curvature depending on membrane tension^36^ and maturation^29^. This process is not understood. In particular, how flat caveolae curve, or how curved caveolae flatten, and which proteins regulate these transitions are unclear. To gain a global view of caveolae density, shapes, and sizes, we analyzed caveolae at the plasma membrane across several common cell types with platinum replica electron microscopy. To visualize caveolae at high resolution, cells were grown on coverslips, unroofed with a light sheering force to expose their inner plasma membranes, and platinum replicas of the cytosolic face of these membranes were generated and imaged by transmission electron microscopy (TEM, Fig. 1A)^37,38^. In these images, caveolae can be identified by their size, round shape, and distinctive striped coat (orange arrows in Fig. 1A). Adipocytes contain substantially more caveolae at their plasma membrane compared to other analyzed cell types (Fig. 1B, caveolae number/μm^2^: MEF 2.6 ± 0.4, adipocytes 14.8 ± 1.4, myoblasts 1.7 ± 0.5, astrocytes 3 ± 0.8, HUVEC 2.4 ± 0.3, HeLa 1.9 ± 0.3). Caveolae had diameters between 40 and 160 nm. In fibroblasts (mouse embryonic fibroblasts, MEFs) and endothelial cells (Human umbilical vein endothelial cells, HUVEC) caveolae diameters were slightly larger compared to other cell types (Fig. 1B, MEF 100.5 ± 2.1 nm, adipocytes 84.9 ± 1.3 nm, myocytes 84.5 ± 1.6 nm, astrocytes 83.2 ± 1.6 nm, HUVEC 98.5 ± 1.6 nm, 80.2 ± 1.2 nm). The caveolae coat contained an average of 4 to 5 visible coat “stripes” with a length of 47.5 ± 1.2 nm (Fig. 1C) and a convoluted spiral-like topology.

**Figure 1:**
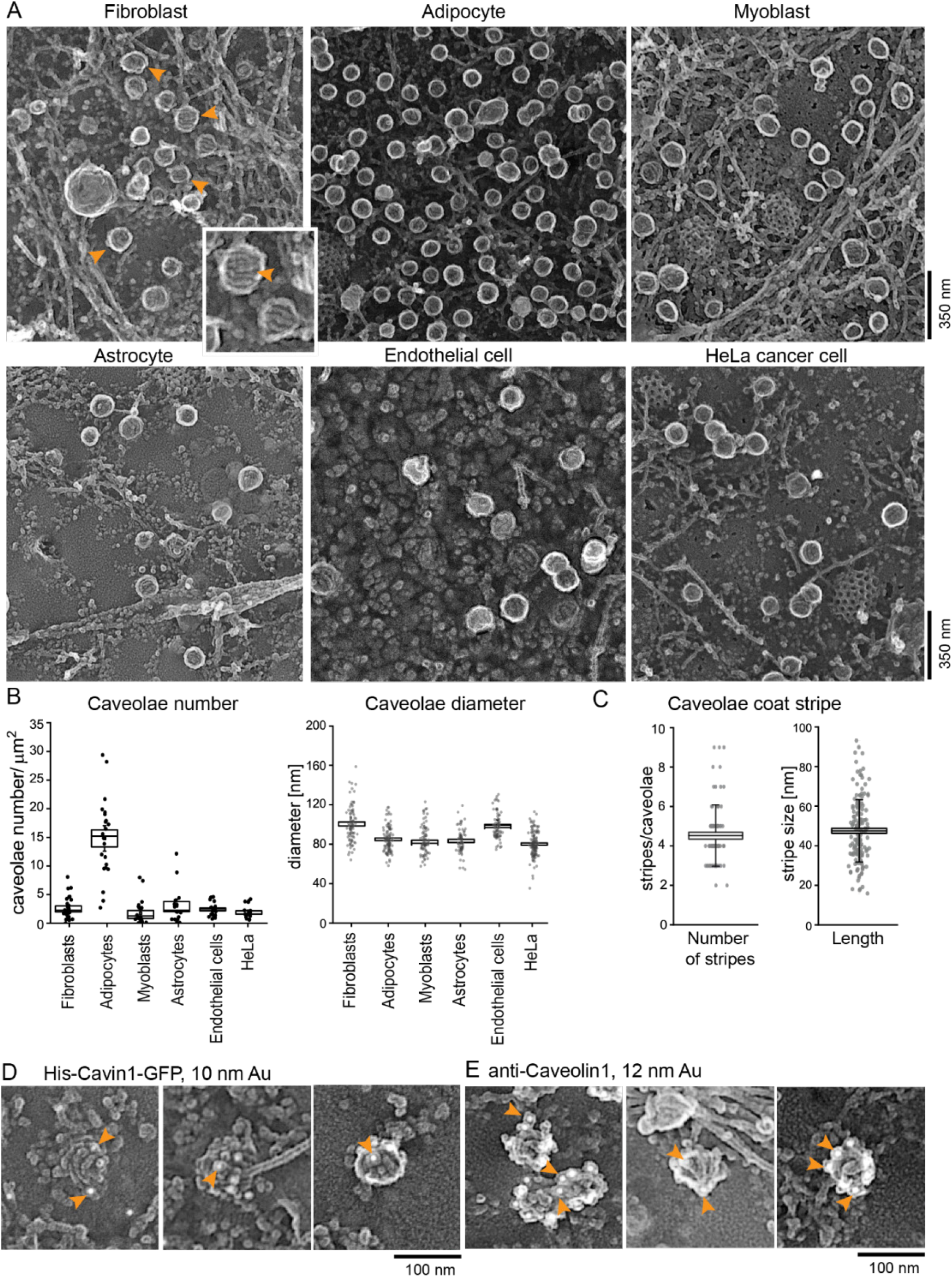
Overview of caveolae at the plasma membrane in various cell types. **(A)** Representative platinum replica transmission electron microscopy (PREM) images of plasma membrane sheets of different cell types. The images are presented in an inverted scale. Orange arrows indicate single caveolae. **(B)** Caveolae number and diameter was measured in all cell types (number: n(fibroblasts) = 26, n(adipocytes) = 22, n(myoblasts) = 22, n(astrocytes) = 17, n(endothelial cells) = 21, n(HeLa) = 19; diameter: n(fibroblasts, MEFs) = 82, n(adipocytes) = 95, n(myoblasts) = 83, n(astrocytes) = 64, n(endothelial cells, HUVEC) = 70, n(HeLa) = 118). **(C)** Number of single coat stripes per caveolae (n = 79), and stripe length in nm (n = 162) of single coat stripes. **(D-E)** Identification of cavin1 (D) or caveolin1 (E) in PREM of MEFs. His-Cavin1-EGFP was expressed in MEFs and 10 nm Ni-NTA nanogold was used to labeled His tags. Caveolin1 was investigated by immunolabeling and 12 nm gold secondary antibody. Orange arrows indicate gold particles. Box plots represent mean ± SE, whiskers show SD, each replicate is depicted.

To verify that these structures were caveolae, the caveolae proteins cavin1 (Fig. 1D) and caveolin1 (Fig. 1E), were labeled with antibody or NTA-linked nanogold particles and imaged with platinum replica EM. Figure 1D shows plasma membranes of MEFs expressing His-tagged cavin1 treated with nickel-NTA labeled gold particles. Caveolin1 was detected by antibody labeling. Both could be detected in all caveolae. Thus, cavin1 and caveolin1 mark morphologically-identifiable caveolae. They also associate with small and disorganized domains with low curvatures.

Next, we quantitively analyzed caveolae curvature in different cell types. Fig. 2A shows example platinum replica EM images of caveolae with a range of morphologies from flat to spherical. The size, packing, and arrangement of the coat varied across structures. Visible stripes could be seen on single caveolae. Flat caveolae exhibit a close packing of the coat and low curvatures (Fig. 2A). Invaginated caveolae showed a similar coat texture, however, the organelle was more curved, forming a bulb shape with a noticeable edge density (white region) relative to the center of the organelle. This edge signal arises from metal accumulated along the side of the organelle when the sample is coated with platinum at an angle. The thicker material blocks passing electrons and appears as a white ring in inverted images. Spherical caveolae have a stronger edge signal with no clear material transitioning from the caveolar body to the surrounding membrane. From these criteria, we classified caveolae into three morphologically classes: flat, bulb, and sphere. Of note, in deeply invaginated caveolae (spheres), the caveolar neck is located under the coat and is therefore hidden when viewed from above^39^. Thus, we cannot ascertain if spherical caveolae are connected or separated from the plasma membrane.

**Figure 2:**
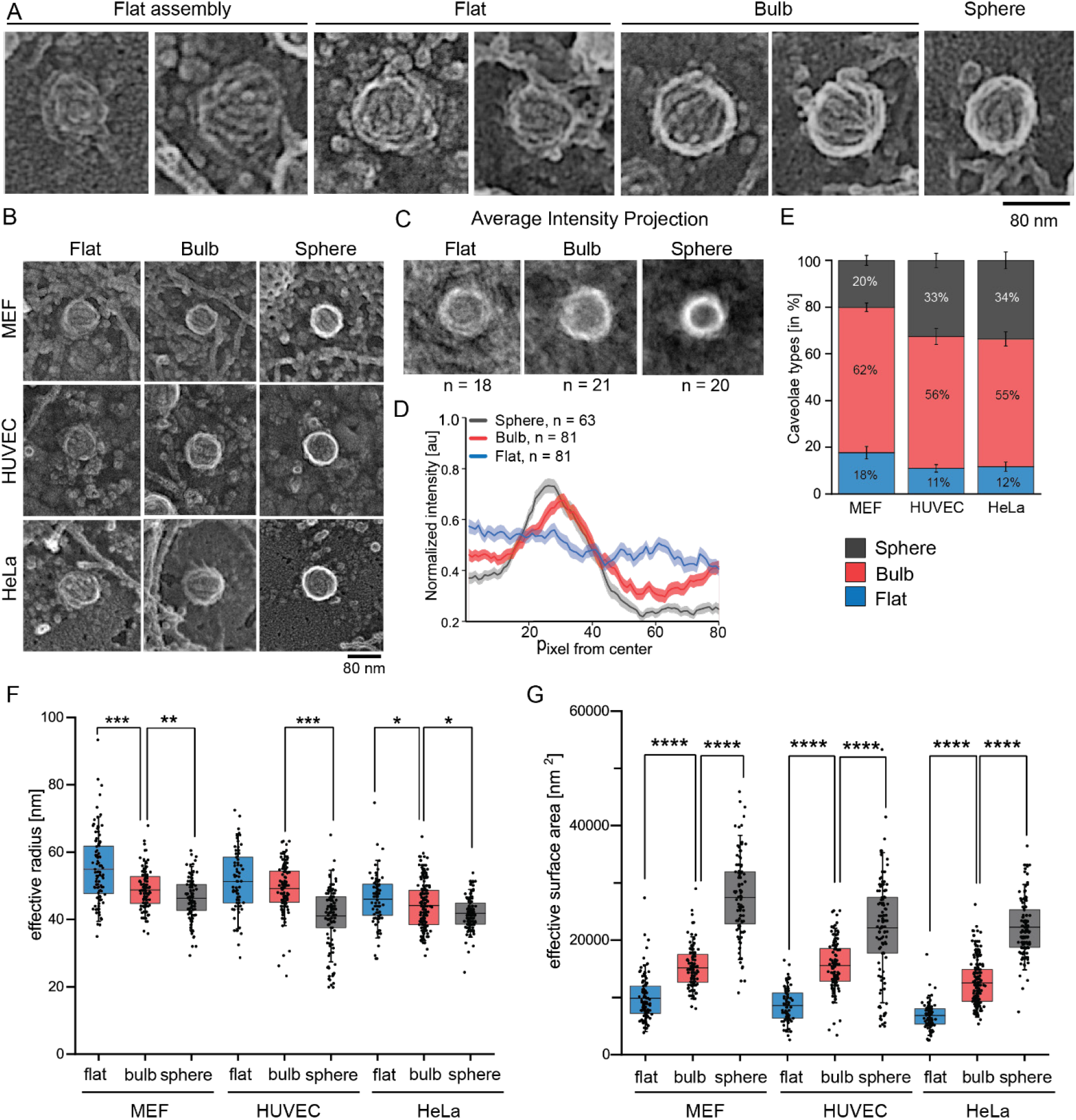
Spherical caveolae show distinct round membrane edges and a smaller size. **(A)** Representative PREM images of caveolae types in which the membrane leaflet can be identified as a dark background and protein by increased electron intensity (grey or white signal, in MEFs). Strong membrane curvature or bending is indicated by a strong white edge signal. **(B)** Representative PREM images of flat, bulb, and spherical caveolae in MEF, HUVEC, and HeLa cells. Scale bar is 80 nm. **(C)** Average intensity projection of 18 flat, 21 bulb, or 20 spherical caveolae in MEFs. **(D)** The intensity plot illustrates normalized electron intensity, mean ± SE from the center of the caveolae domain to its edge (n(flat) = 63/4 cells, n(bulb) = 81/4 cells, n(sphere) = 81/4 cells). **(E)** Distribution of caveolae types in MEF, HUVEC and HeLa (n(MEF) = 12, n(HUVEC) = 13, n(HeLa) = 12). Bar plot shows mean ± SE. **(F)** Effective radius of caveolae was calculated with the assumption of round caveolae membrane domains (MEF: n(flat) = 100, n(bulb) = 113, n(sphere) = 96; HUVEC: n(flat) = 67, n(bulb) = 108, n(sphere) = 113; HeLa: n(flat) = 76, n(bulb) = 177, n(sphere) = 133). **(G)** Effective surface area of flat (based on circle: A = πr^2^), bulb (based on hemisphere: A = 2πr^2^), and spherical caveolae (based on sphere: A = 4πr^2^; MEF: n(flat) = 100, n(bulb) = 113, n(sphere) = 96; HUVEC: n(flat) = 67, n(bulb) = 108, n(sphere) = 113; HeLa: n(flat) = 76, n(bulb) = 177, n(sphere) = 133). Box plots represent mean ± SE, whiskers illustrate SD, each replicate is depicted. Normal distributed groups were analyzed by t-test, not normally distributed values with Mann Whitney test, *P < 0.05, **P < 0.01, ***P < 0.0001, ****P < 0.00001.

Platinum replicas of unroofed MEFs, HUVEC, and HeLa cell plasma membranes were generated, imaged, and segmented for all visible caveolae. Fig. 2B shows caveolae shapes (flat, bulb, sphere) across the three cell types. The average intensity projections of flat, bulb, or sphere caveolae illustrates the average measurable differences in curvature and sizes across the caveolae types (Fig. 2C). When line scans were normalized and plotted from the center of the individual organelles towards their edges, different caveolae types could be classified by their unique profiles (Fig. 2D). Flat caveolae exhibited minimal intensity difference between the center, edge, and outside of the organelle image (blue graph in Fig. 2D). Yet, a clear edge signal was detectable in bulb shaped caveolae (Fig. 2B, C) that was reflected in an intensity maximum in Fig. 2D (red graph). Sphere-shaped caveolae showed a steeper slope and larger intensity difference that was shifted towards the center of the organelle (black graph in Fig. 2D). In support of these measurements, electron tomograms of flat caveolae showed a lower height and lower curvature compared to bulb or spherical caveolae (Fig. S1, Video S1-4).

Fig. 2B shows that in all three cell types, flat, bulb, and sphere caveolae were detected (Fig. 2B, E). MEFs contained slightly more flat and bulbed shapes compared to HUVEC and HeLa cells (Fig. 2E). Flat caveolae were wider than spherical caveolae, indicating that coat bending decreases the diameter of caveolae (Fig. 2F). Estimated surface areas could be calculated based on the radii^40^ where flat caveolae were assumed as circles, bulbs as hemispheres, and spheres as spheres. Given these changes, the calculated surface area of the organelle increased during invagination (Fig. 2G). Thus, caveolae capture additional membrane and swell into the cell, decreasing the cell’s exposed surface area when the coat assembles and bends.

### STED microscopy of single caveola in plasma membrane sheets

To localize caveolae-related proteins on single caveolae at the nanoscale, we developed a two-color super-resolution fluorescence microscopy (stimulated emission depletion, STED) method. First, MEF plasma membrane sheets were immuno-stained against caveolin1 (a marker for caveolae, Fig. 3A). As illustrated in Fig. 3B, sufficient lateral resolution could be achieved with the STED dye Atto647N (see also Fig. S2A, B) to visualize single caveolin1 spots. Next, two-color STED was used to localize cavin1 in relation to Caveolin1 (Fig. 3C, D). To image Cavin, Cavin1-EGFP was expressed in MEFs and labelled with Atto647-GFP-nanobodies. Caveolin1 was detected by immunolabelling with anti-caveolin antibodies and Alexa594-secondary antibodies (Fig. 3C, D). Compared to the caveolae diameter measured by PREM (101 ± 2 nm, Fig. 1B), the size of caveolin1-positive STED spots labeled with Atto647N showed no substantial size differences across the averages with either GFP-nanobody (114 ± 6 nm) or antibody immunolabeling (108 ± 7 nm, Fig. S2C, D). Alexa594 labeling suggested a slightly larger size of 138 ± 7 nm.

**Figure 3:**
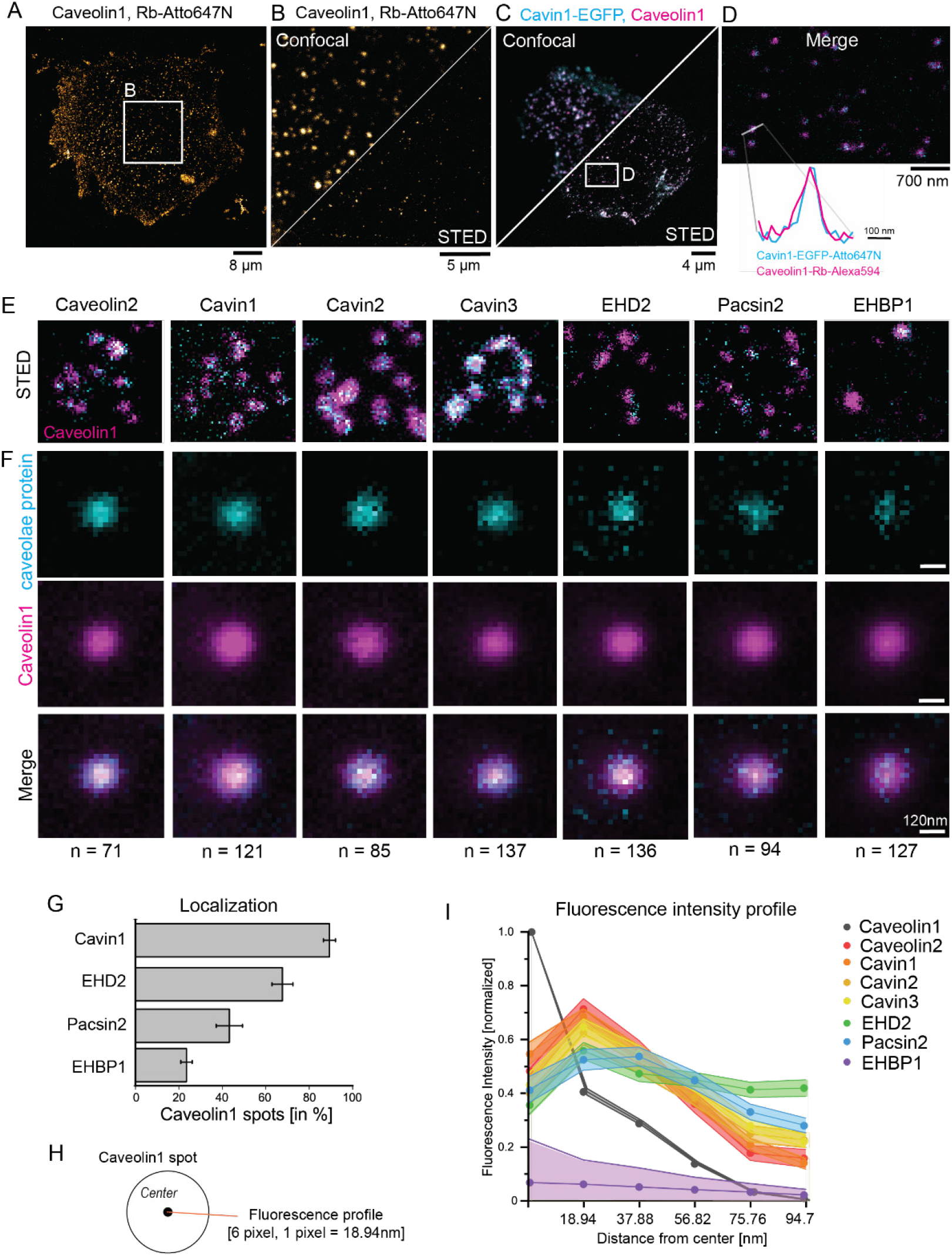
Stimulated emission depletion microscopy (STED) shows specific protein profiles for caveolae. **(A)** Confocal image of MEF plasma membrane sheet immunolabelled with an antibody against caveolin1 and a secondary anti-rabbit antibody tagged with Atto647N (Rb-Atto647N). **(B)** Enlarged selection from (A) shows confocal and STED image of endogenous caveolin1 in MEFs. **(C-D)** Confocal and STED image of cavin1-EGFP expressing MEFs immunolabelled with caveolin1 antibody (secondary antibody dye Alexa594, magenta). Cavin1 was tagged by GFP nanobody labelled with Atto647N (blue). (D) shows enlarged STED image section. STED fluorescence profile illustrates cavin1 localization to a caveolin1 spot. **(E)** Representative STED images of caveolae proteins (cyan) and caveolin1 antibody labeling (magenta, secondary antibody tagged with Alexa594) in plasma membrane sheets from MEFs. The individual caveolae proteins were expressed with EGFP tags and labelled with GFP nanobody-Atto647N. **(F)** Normalized average STED fluorescence intensity projection of automatically detected caveolin1 spots (magenta) and the corresponding co-labeled caveolae proteins (cyan). Lower panel shows both channels as merged image and the total number of caveolin1 spots is indicated. Scale bar represents 120 nm. **(G)** Percentage of caveolin1 spots that showed localization of either Cavin1, EHBP1, EHD2 or Pacsin2. Bar plot indicates mean ± SE (n(Cavin1) = 1135/8 cells, n(EHBP1) = 1452/11 cells, n(EHD2) = 949/8 cells, n(Pacsin2) = 1037/9 cells). **(H)** The STED fluorescence plot profile for the individual caveolae proteins was analyzed from the center of the caveolin1 spot to the edge. Pixel size in STED images was 18.94 nm, based on the estimated caveolae diameter of 100 nm (radius = 50 nm), the edge of caveolae can be assumed between 3-4 pixel from the center (56-75nm). **(I)** Fluorescence plot profiles from the center of caveolin1 spots accordingly to (H). Line graph indicates mean ± SE for each pixel and caveolae protein (n(Caveolin1) = 121, n(Caveolin2) = 71, n(Cavin1) = 121, n(Cavin2) = 85, n(Cavin3) = 137, n(EHD2) = 136, n(Pacsin2) = 94, n(EHBP1) = 127).

Next, we investigated the localization of the coat proteins caveolin2 and cavins in relation to caveolin1 with two-color STED (Fig. 3E, F). EGFP-tagged caveolin2, cavin1, 2, or 3 were expressed and labelled with a GFP nanobody-Atto647N probe. Caveolin1 was immunolabelled and detected with Alexa594. As expected, cavin coat proteins strongly colocalized with caveolin1 (Fig. 3E). In contrast, the caveolar regulatory proteins EHD2 and pacsin2, as well as EHBP1, exhibited a more punctate localization at caveolin1 spots (Fig. 3E). This produced a weaker average fluorescence signal relative to the background when many spots were aligned and averaged (Fig. 3F). Cavin1 was detected at more than 90% of all caveolin1 spots. Pacsin2 and EHBP1 were observed at 40% or 20% of all caveolin1 spots, respectively (Fig. 3G). These data suggest that both EHBP1 and pacsin2 localize to a subset of caveolae at any given time. Analyzing the average fluorescence profiles of caveolae-associated proteins (Fig. 3H), all caveolae coat proteins shared a similar distribution at STED resolutions compared to caveolin1 (Fig. 3I). EHD2 and pacsin2 exhibited a slightly extended shape. In summary, STED microscopy can visualize multiple proteins at single caveolae at the plasma membrane. We hypothesized that pacsin2 and EHBP1 may associate with a specific caveolar shape.

### STED Pt replica CLEM reveals cavin localization to flat and curved caveolae types

To test this hypothesis, we directly correlated STED images to Pt replica of the same samples with a correlative light and electron microscopy method (CLEM). Here, caveolae protein localizations could be directly compared to the curvature of the organelle and the surrounding cell membrane (STED-CLEM, Fig. S3 overview of correlated MEF). First, MEFs expressing caveolin1-EGFP or caveolin2-EGFP were labeled with a GFP nanobody-Atto647N and imaged (Fig. 4A, B). This approach made it possible to detect caveolin1 accumulation at all stages of caveolae formation (right panel Fig. 4A, diffused STED signal). As the caveolae coat is formed, the caveolin1 STED signals mark morphologically and EM-identifiable caveolae. As expected, caveolin1 was found in all caveolae types. Caveolin2 showed a similar behavior (Fig. 4B). Quantitative analysis of the fluorescence associated with the EM segmented caveolae regions indicated that caveolin1 and 2 profiles for flat or spherical caveolae had similar distributions (Fig. 4C, edge of caveolae indicated in green dashed line).

**Figure 4:**
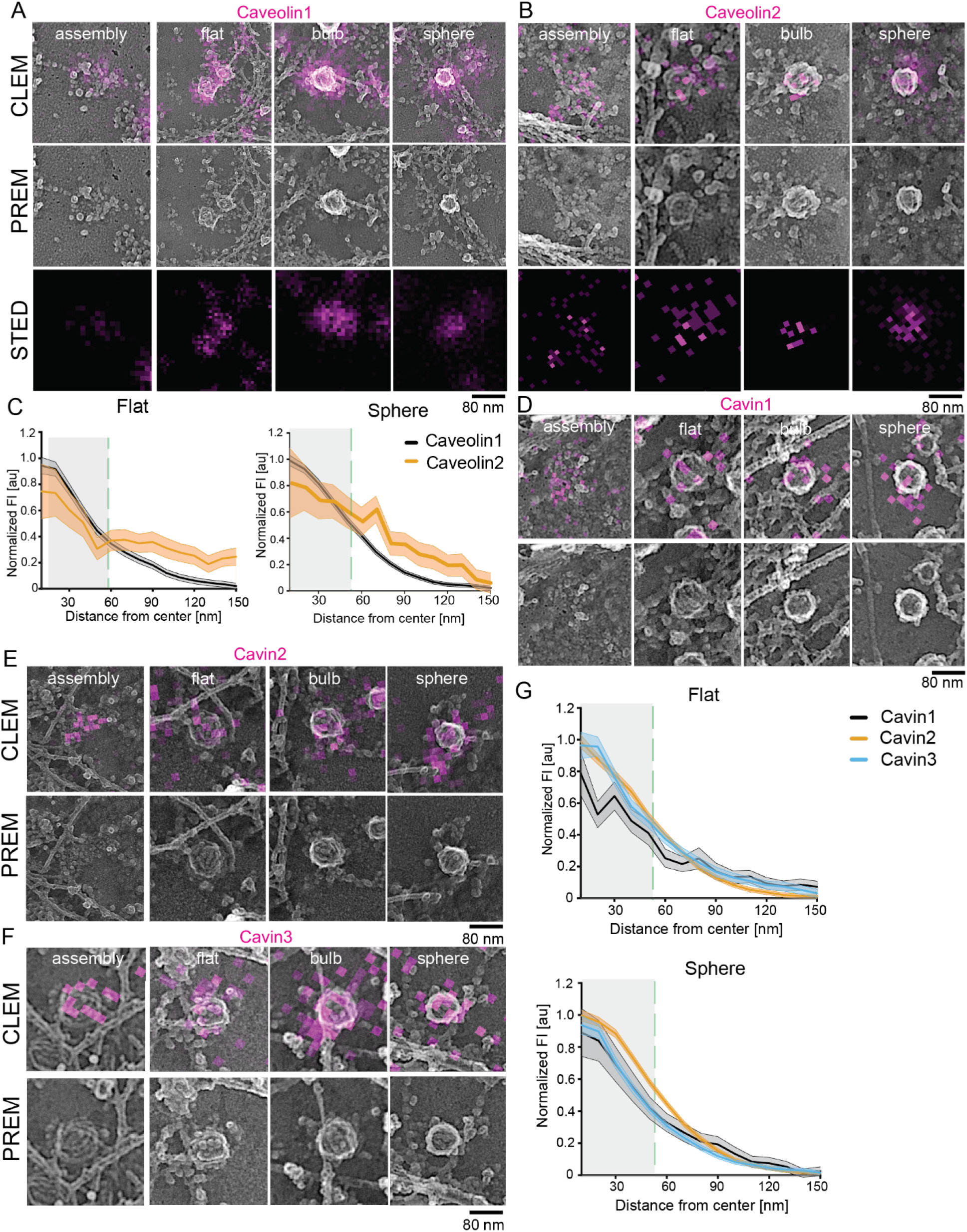
STED-CLEM revealed the localization of cavin proteins to flat, bulb and spherical caveolae. **(A-B)** Representative CLEM images for MEFs expressing either caveolin1-EGFP (A) or caveolin2-EGFP (B) which were labelled with GFP nanobody-Atto647N and imaged with STED and PREM. **(C)** Quantitative analysis of flat or spherical caveolae STED fluorescence profiles from the center of caveolae to their edges (indicated by green dashed line, line graphs show mean ± SE; caveolin1: n(flat) = 145, n(sphere) = 178; caveolin2: n(flat) = 140, n(sphere) = 474). **(D-F)** Representative CLEM images for MEFs expressing either cavin1-EGFP (D), cavin2-EGFP (E), or cavin3-EGFP (F) labelled with GFP nanobody-Atto647N and investigated by STED followed by PREM. **(G)** Quantitative analysis of flat or spherical caveolae by STED fluorescence profiles from the center of caveolae to their edges (indicated by green dashed line; line graphs show mean ± SE; cavin1: n(flat) = 59, n(sphere) = 272; cavin2: n(flat) = 231, n(sphere) =223; cavin3: n(flat) = 198, n(sphere) = 191).

We next imaged cavin1-3 (Fig. 4D-G). We detected all three cavin isoforms in immature caveolae where the membrane domains were small and disorganized (Fig. 4D-F, subpanel ‘assembly’). Importantly, all cavins were found associated with flat caveolae, similar to data from immunogold labeling (Fig. 1C). Analysis across the entire population of labeled caveolae (n= 488) indicated that cavin1-3 localized to spherical, bulb, and flat caveolae (Fig. 4G). To further confirm this, we imaged endogenous cavin1 in Pt replicas of endothelial cells after hypo-osmotic shock (Fig. S4). As shown previously^33^, hypo-osmotic shock in HUVEC resulted in an increase in flat caveolae (Fig. S4A-C). When cavin1 was immuno-labelled, again we detected cavin1 on all shapes (Fig. S4D, E). Quantitative analysis of 372 flat caveolae (in 8 different STED-CLEM regions) showed that 84% were cavin1 positive (Fig. S4F). This is contrary to current models of cavin behavior, where the cavin coat is proposed to be disassembled and lost when caveolae are flat.

### The caveolae regulatory proteins EHD2 and pacsin2 accumulate at flat caveolae

Next, we investigated the localization of EHD2 and pacsin2. First, EHD2-EGFP expressing MEFs were imaged by STED-CLEM (Fig. 5A). Previous work has suggested that EHD2 localizes to the neck of caveolae^18^. Correlated STED data, however, showed that EHD2 was also detected at flat caveolae and immature caveolar domains (Fig. 5A, B). Interestingly, the EHD2 fluorescence signal which is diffuse in flat structures was concentrated around 20 nanometers from the center of bulb and spherical caveolae (as illustrated in the fluorescence plot profile in fig. 5B). This suggests that EHD2 molecules reposition to the caveola neck during curvature acquisition, similar to the dynamics of the related protein dynamin at clathrin-coated pits. Because overexpression of EHD2 results in and increase in curved and immobile caveolae, we also imaged endogenous EHD2 with antibodies. Fig. 5C shows representative antibody-stained CLEM images of flat caveolae coated with EHD2. Similar to EHD2-EGFP expressing cells, quantitative analysis indicated a distinct signal of EHD2 at both flat and curved caveolae (Fig. S5C).

**Figure 5:**
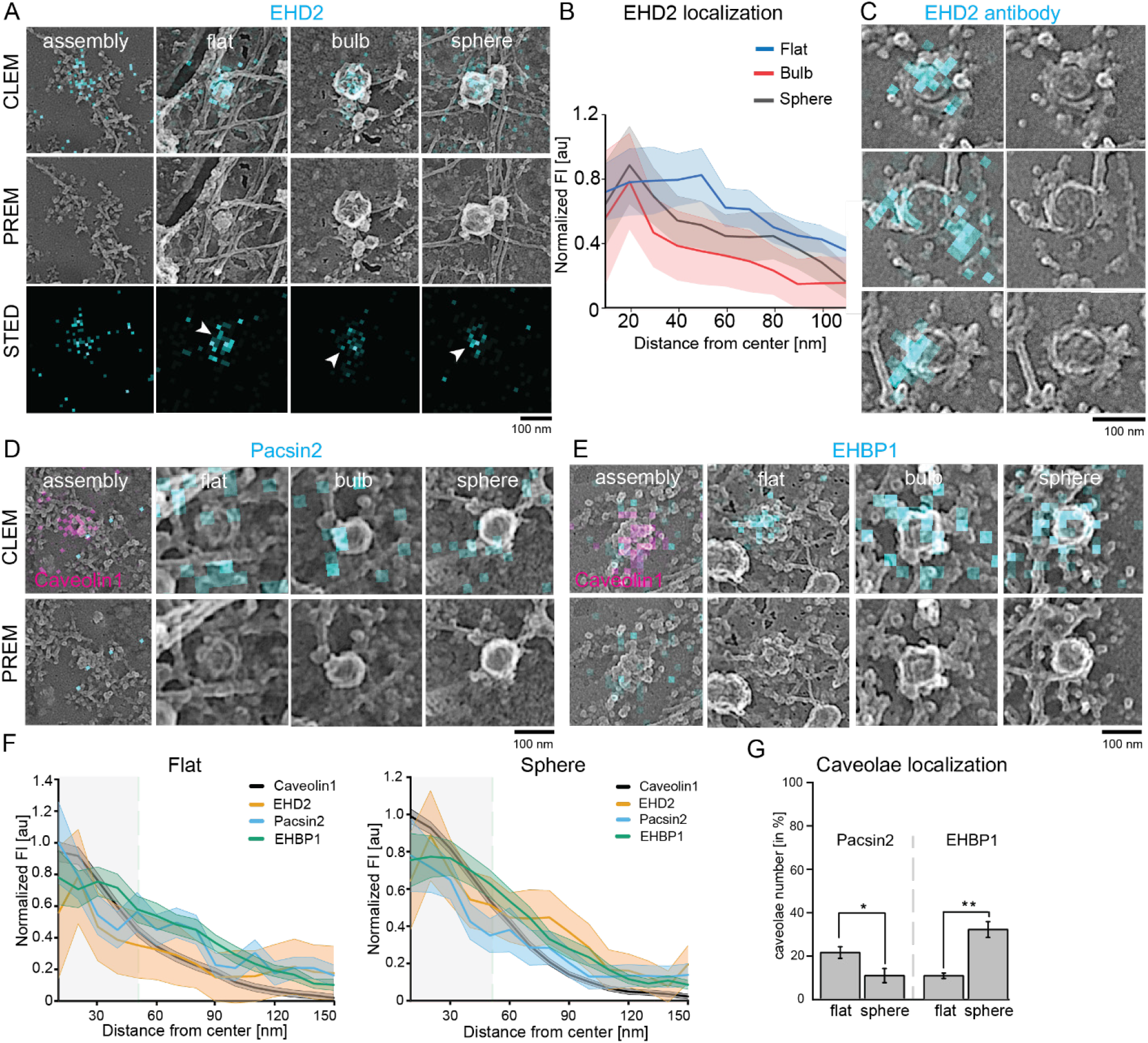
Spatially distribution of EHD2, pacsin2 and EHBP1. **(A)** Representative STED-CLEM images for MEFs expressing EHD2-EGFP (A, labelled with GFP nanobody-Atto647N) or white arrows indicate accumulation of EHD2 around the caveolae center. **(B)** EHD2 STED fluorescence profile from the center of caveolae to its edges obtained from STED-CLEM images (A). Each individual caveolae type is depicted (graph shows mean ± SE, EHD2: n(flat) = 72, n(bulb) = 50, n(sphere) = 163). **(C)** STED-CLEM showing endogenous EHD2 antibody staining at flat caveolae (secondary antibody tagged with Atto647N). **(D-E)** Representative CLEM images for MEFs expressing either pacsin2-EGFP (C) or EHBP1-EGFP (D) that were labelled with GFP nanobody-Atto647N (illustrated in cyan, caveolin1 STED signal in magenta). **(F)** Quantitative analysis of flat or spherical caveolae by STED fluorescence profiles from the center of caveolae to their edges (indicated by green dashed line, line graphs show mean ± SE, caveolin1: n(flat) = 145, n(sphere) = 178; EHD2: n(flat) = 72, n(sphere) = 163; pacsin2: n(flat) = 103, n(sphere) = 77; EHBP1: n(flat) = 70, n(sphere) = 79). **(G)** Percentage of flat or spherical caveolae that were targeted by either pacsin2 or EHBP1. Bar plot indicates mean ± SE, n (pacsin2) = 3 cells, n(EHBP1) = 4 cells. Scale bar is 100 nm.

Only a sub-population (42%) of caveolin1 spots were positive for pacsin2 in the previous STED experiment (Fig. 3G). To identify which caveolae sub-type is enriched in pacsin2, MEFs expressing pacsin2-EGFP were analyzed by STED-CLEM (Fig. 5D, F). Flat caveolae displayed elevated pacsin2 occupancy compared to spherical caveolae (Fig. 5G). These observations were verified by antibody staining for endogenous pacsin2, which showed a similar preferred association with flat structures (Fig. S5A, C).

### EHBP1 localizes to curved caveolae

The actin and EHD2 binding protein EHBP1 associates with caveolae^16^, however its location is unclear. Therefore, STED-CLEM of MEFs overexpressing EHBP1-EGFP was used to image EHBP1. Fig. 5E shows representative CLEM images for EHBP1. Quantitative analysis showed that EHBP1 accumulates at spherical caveolae rather than flat domains (Fig. 5G). Antibody staining for EHBP1 confirmed this distribution (Fig. S5B).

In summary, at nascent caveolae domains, cavin proteins accumulate with caveolins when the characteristic well-defined coat is not yet clearly formed. Cavin proteins remain on flat caveolae. EHD2 is present at all caveolae domains despite the lack of a caveolae neck. Pacsin2 primarily localizes to flat caveolae, and EHBP1 is enriched in spherical caveolae.

### Caveolae regulatory proteins restrain caveolae in a less curved state

Next, we tested how reduced levels of EHD2, pacsin2, or EHBP1 effect caveolae shape and density (Fig. 6A, Western Blot verification Fig. S6). MEFs lacking EHD2^17^ exhibited no change in the number of caveolae (Fig. 6A-C), however substantially more spherical caveolae were observed compared to wild-type MEFs (Fig. 6D). siRNA based knockdown of pacsin2 resulted in an increase in flat caveolae (40% vs. 24% in wild-type), while reducing the percentage of spherical caveolae (14% from 25% in wild-type, Fig. 6A-D). Knockdown of EHBP1 did not alter caveolae number or proportion of types (Fig. 6D). Interestingly, the loss of all three proteins (double knockdown of Pacsin2 and EHBP1 in EHD2 lacking cells, triple KO/KD, see also Western Blot in Fig. S6) substantially reduced the total number of caveolae (Fig. 6C). In particular, flat and bulb caveolae percentages were reduced in this condition compared to wild-type cells. Surprisingly, the triple KO/KD MEFs showed more spherical caveolae in comparison to wild-type MEFs (Fig. 6D). 53% of all caveolae were spheres in triple KO/KD MEFs compared to 25% in wild-type MEFs. Notably, caveolin1 and cavin1 protein levels were also reduced in triple KO/KD MEFs (Western Blot Fig. S6C). Size measurements for the individual caveolae showed that the loss of EHD2, pacsin2, or EHBP1 results in smaller caveolae with the exception of flat caveolae in Pacsin2 knockdown MEFs (Fig. 6E).

**Figure 6:**
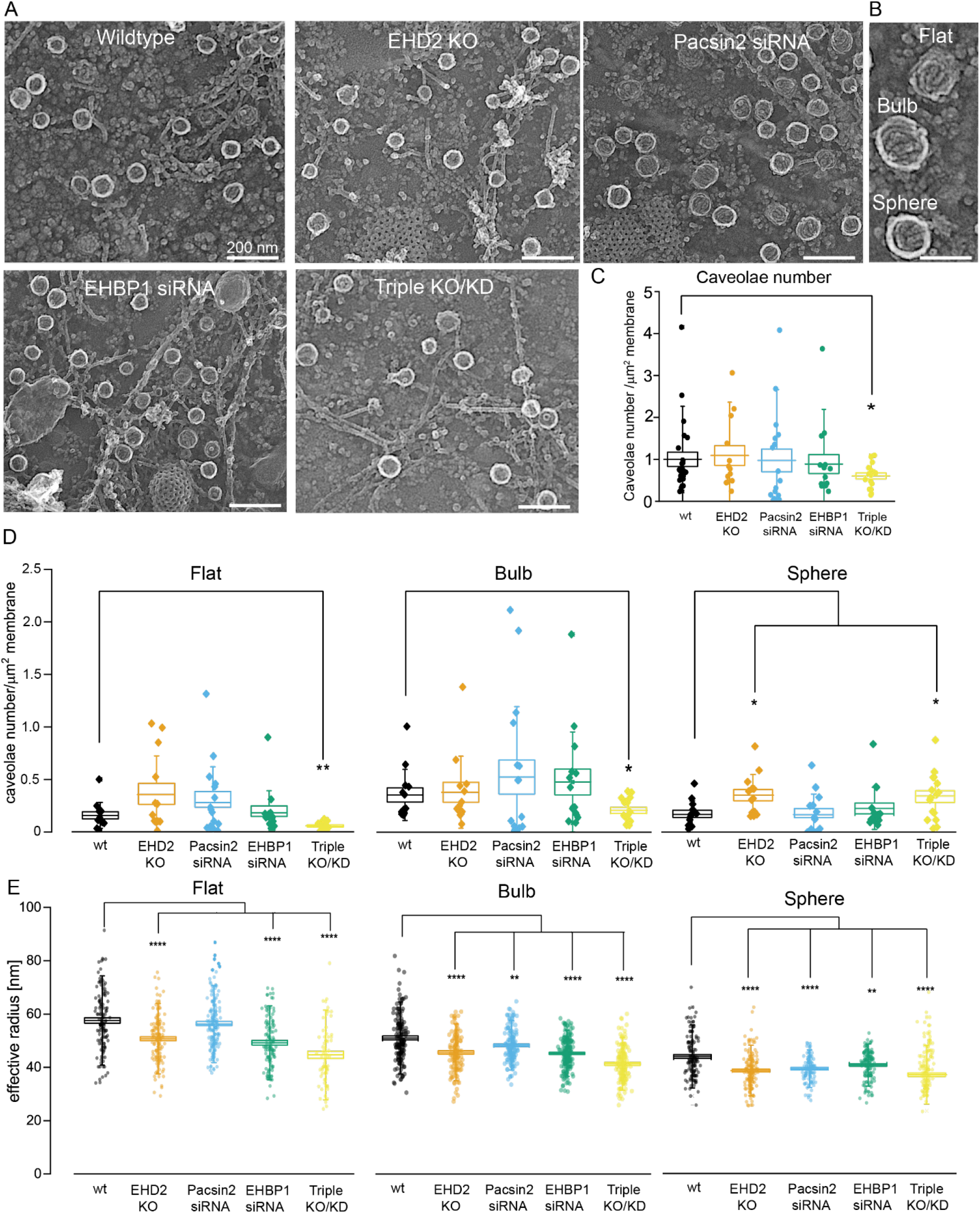
EHD2, pacsin2 and EHBP1 stabilize caveolae at the plasma membrane. **(A)** Representative PREM example images of wildtype MEFs (wt), EHD2 knockout MEFs (KO), pacsin2 siRNA or EHBP1 siRNA treated MEFs. Triple knockout/knockdown (KO/KD) indicates EHD2 KO MEFs that were treated with pacsin2 and EHBP1 siRNA. Scale bar is 200 nm. **(B)** Example PREM image of flat, bulb and spherical caveolae. **(C)** In PREM images the total caveolae number at the plasma membrane in wt, EHD2 KO, pacsin2 siRNA treated, EHBP1 siRNA treated, or triple KO/KD MEFs was measured. Box plot indicates mean ± SE, whiskers show SD, each replicate is depicted (n(wt) = 25, n(EHD2 KO) = 13, n(pacsin2 siRNA) = 16, n(EHBP1) = 15, n(Triple KO/KD) = 16). **(D)** Number of caveolae types in wt, EHD2 KO, pacsin2 siRNA, EHBP1 siRNA or triple KO/KOD MEFs. Box plot indicates mean ± SE, whiskers show SD, each replicate is illustrated (n(wt) = 13, n(EHD2 KO) = 13, n(pacsin2 siRNA) = 17, n(EHBP1) = 15, n(Triple KO/KD) = 16). **(E)** Caveolae radius (round caveolae domains were assumed) of flat, bulb or spherical caveolae in wt, EHD2 KO, pacsin2 siRNA, EHBP1 siRNA or triple KO/KOD MEFs. Box plot indicates mean ± SE, whiskers show SD, each replicate is illustrated (flat: n(wt) = 127, n(EHD2 KO) = 139, n(pacsin2 siRNA) = 163, n(EHBP1) = 123, n(Triple KO/KD) = 69; bulb: n(wt) = 148, n(EHD2 KO) = 130, n(pacsin2 siRNA) = 146, n(EHBP1) = 177, n(Triple KO/KD) = 145; sphere: n(wt) = 129, n(EHD2 KO) = 157, n(pacsin2 siRNA) = 106, n(EHBP1) = 142, n(Triple KO/KD) = 133). **(F)** Statistical significance was measured by t-Test in normally distributed data sets, otherwise the nonparametric Mann Whitney test was applied. *P < 0.05, **P < 0.01, ***P < 0.0001, ****P < 0.00001.

### Dynamin does not localize to caveolae

Dynamin has been implicated in caveolae endocytosis^19^. Caveolae mobility and endocytosis are inhibited in cells expressing the dynamin mutant K44A (Dyn-K44A)^17,23,41^. However, previous studies largely failed to strongly localize dynamin at caveolae. Therefore, we mapped dynamin at individual caveolae. First, STED was used to detect dynamin2-EGFP (the major isoform in MEFs^42^) at the plasma membrane (Fig. 7A). Surprisingly, no substantial co-localization of dynamin2 (Fig. 7A, cyan) and caveolin1 (Fig. 7A, magenta) was observed. In contrast, Dynamin2 was commonly and strongly detected at clathrin sites in these cells (Fig. 7A, yellow). The Dyn-K44A mutant has been proposed to accumulate at caveola necks^13,22,43^. In contrast, we rarely observed colocalization of Dyn-K44A and caveolin1 above background levels (Fig. 7A, lower panel). Quantitative analysis showed that dynamin2 or Dyn-K44A localized to only 8.9 ± 1.1% or 18.2 ± 2.4% of caveolae. Dynamin mutants showed a modest increase over background (Fig. 7B, C). Therefore, Dyn-K44A was used to study dynamin in STED-CLEM (Fig. 7D). As expected, dynamin was strongly localized to clathrin-coated sites (Fig. 7D -b). However, dynamin did not strongly localize with caveolae (Fig. 7D -c). To complement these observations from transfected cells, MEFs were immuno-stained against endogenous dynamins. Again, no localization of dynamin at caveolae was detected (Fig. S7). Furthermore, STORM-CLEM of dynamin2 in HeLa and SK-MEL-2 cells^38^ did not reveal strong association of dynamin with caveolae (Fig. S8, dynamin-GFP and endogenous dynamin). Additionally, Dyn-K44A STORM-CLEM in SK-MEL-2 cells did not show robust accumulation at caveolae (Fig. S8E, F). Commonly, dynamin2 bound to clathrin would sit near caveolae. This close association between the two organelles might lead to false-positive co-localizations and may explain past suggestions of co-localizations between caveolae and dynamin (Fig. S8B). These false positives would be invisible without CLEM or three color super-resolution imaging of dynamin, clathrin, and caveolae.

**Figure 7:**
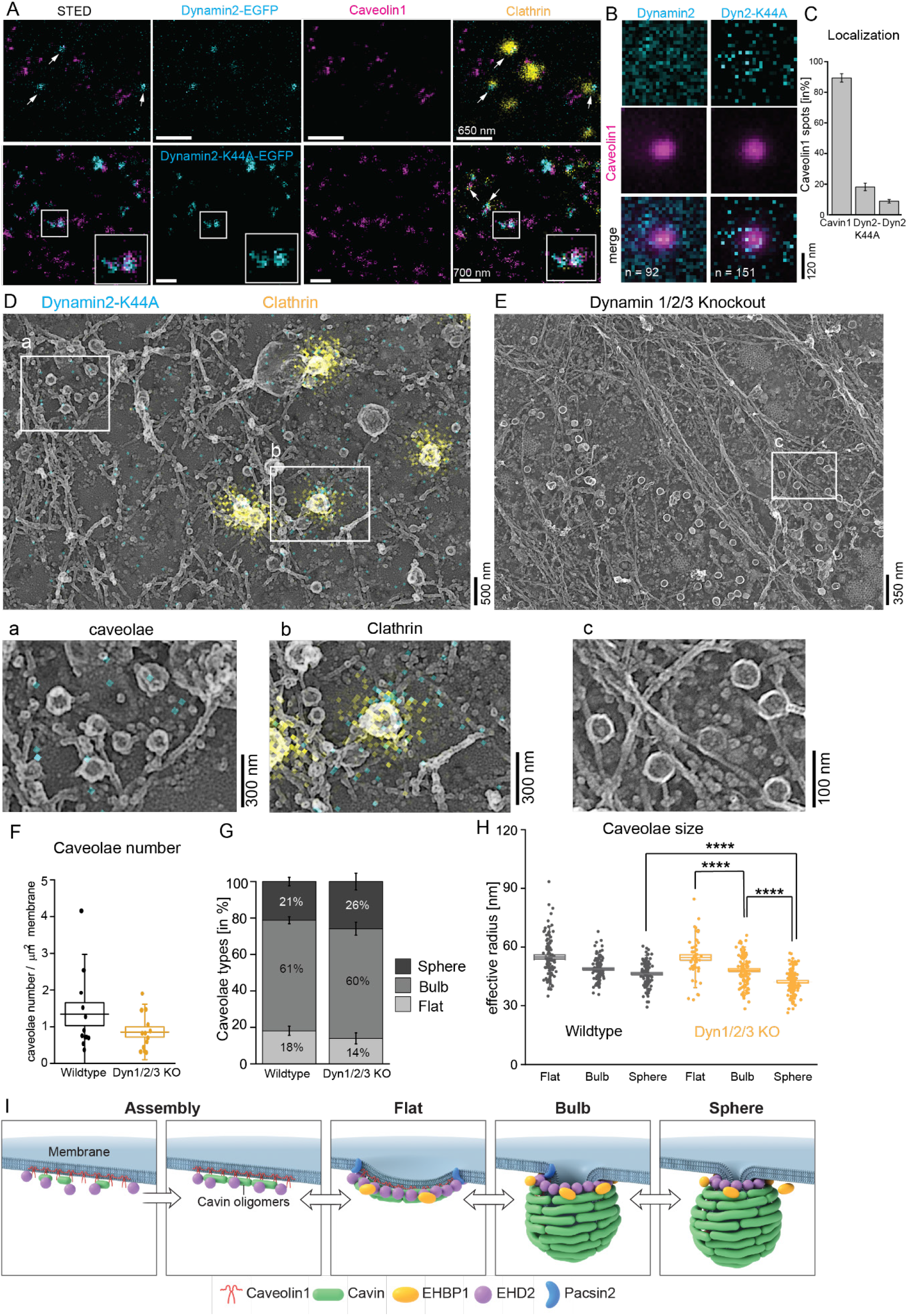
Dynamin is not localized to caveolae. **(A)** Representative STED images of MEF membrane sheets expressing dynamin2-EGFP or dynamin2-K44A (in cyan), together with caveolin1 (magenta) and clathrin (yellow) immuno-staining. White arrows indicate dynamin localization. **(B)** Quantitative analysis of dynamin2 or dynamin2-K44A localization to caveolin1 spots in STED images illustrated in average STED fluorescence intensity projections (n(dynamin2) = 92, n(Dyn2-K44A) = 151). **(C)** Percentage of caveolin1 spots that were also stained for cavin1, Dyn2-K44A or dynamin2. Bar plot indicates mean ± SE (n(cavin1) = 1135/8 cells, n(Dyn2-K44A) = 1283/11 cells, n(dynamin) = 1032/8 cells). **(D)** Representative STED-CLEM image showing Dyn2-K44A (cyan) and clathrin (yellow) on Pt replica TEM image. Increased panel a illustrates caveolae, panel b shows clathrin vesicle. **(E)** Representative PREM image of membrane sheet obtained from dynamin triple knockout (Dynamin1/2/3) MEFs. Panel c illustrates increased membrane area covered with caveolae. **(F)** Total caveolae number at the plasma membrane in wild-type and dynamin triple knockout MEFs. Box plot shows mean ± SE, whiskers illustrate SD (n(wt) = 39, n(Dyn 1/2/3 KO) = 13). **(G)** Percentage of caveolae types observed in plasma membrane sheets of wild-type and dynamin triple knockout MEFs. Bar graph indicates mean ± SE (n(wt) = 39, n(Dyn 1/2/3 KO) = 13). **(H)** Radius of individual caveolae types (round caveolae domain was assumed), box plot shows mean ± SE, whiskers illustrate SD (wt: n(flat) = 100, n(bulb) = 113, n(sphere) = 96; Dyn 1/2/3 KO: n(flat) = 50, n(bulb) = 110, n(sphere) = 121). Significant difference was tested by Mann Whitney test, **** P < 0.0001). **(I)** Schematic model of caveolae formation and bending of the caveolae core complex.

Next, we tested if there was a morphological change to caveolae when dynamin was knocked out. Specifically, we investigated caveolae number and shape in MEFs lacking all three dynamins. If dynamin is involved in caveolae membrane scission, loss of dynamin should increase caveolae number. PREM of dynamin triple knockout (dynamin1/2/3 knockout^44^, Fig. S9 Western Blot validation) cells showed no obvious changes in caveolae shape or density (Fig. 7E). Quantitative analysis of caveolae number at the plasma membrane did not reveal a significant difference compared to wild-type cells (Fig. 7F). Furthermore, the percentage of flat, bulb, or spherical caveolae was unchanged in dynamin-lacking cells (Fig. 7G). Spherical caveolae showed a slight reduction in size compared to wild-type MEFs (Fig. 7H). The previously-observed size decreases from flat to spherical caveolae (Fig. 2F) was also detected. In summary, we find no strong evidence that dynamin localizes to caveolae or has a mechanistic role in these organelles. These data indicate that dynamin’s previously-reported effects on caveolae could be indirect, very short lived, due to effects on clathrin-mediated endocytosis, general membrane traffic, membrane tension, or the actin cytoskeleton. Future work is needed.

## Discussion

Caveolae are one of the most common organelles found at the plasma membrane of many human cell types. It is still unclear, however, how caveolae assemble, change shape, and are captured into the cell. We analyzed the nanoscale architecture of caveolae marked with key proteins. We classified caveolae into three sub-types according to curvature: flat, bulb, or sphere. In the different cell lines we imaged the percentage of flat, bulb or spherical caveolae was similar. The majority of caveolae are bulb shaped (55-62%). A smaller number are flat (11-18%). Likely, prior to endocytosis, caveolae transition into spheres (20-34%), forming the highly-constricted neck needed for membrane scission. Spherical caveolae likely spend a short amount of time in this state at the plasma membrane before transport into the cytosol. The smaller number of spheres compared to bulb shaped caveolae may reflect the reported limited number of mobile, endocytic caveolae^45^.

We measured a reduction in the radii of caveolae across flat, curved, and spherical shapes. A reduced radii in spherical caveolae has also been observed after extracellular lipid treatment^46^. While the radii decreased, the calculated surface area of individual caveolae increased as caveolae curve, indicating that bending captures excess plasma membrane. Thus, during caveolae curvature, a cell will capture and reduce the exposed plasma membrane surface. Indeed, previous data showed that lipid accumulation in caveolae bulbs can prime invaginations for endocytosis^46^. Possibly, new lipids are required for this process^45^. This could be an important mechanism for lipid uptake.

How do proteins drive membrane curvature at caveolae? First, caveolae, regardless of shape, contained three major proteins (caveolin, cavin, EHD2). Unlike past models, cavins1-3 were found at flat caveolae. This was also true after treating cells with an mild osmotic shock to flatten caveolae. Here, cavin1 remains associated. In past studies, cavins have been proposed to localize only to curved caveolae and were lost when caveolae flatten^3,33^. The loss of cavin was proposed to drive flattening. What is the difference between our studies and past work? Previously, caveolae flattening was induced by very strong and prolonged osmotic shock or cellular stress. These might trigger extreme changes such as an increase in membrane tension, changes in actin polymerization, post-translational modifications, and metabolic changes that might perturb cavin associations. Natural flattening or mild osmotic shock-induced flattening could be a more dynamic and reversible where proteins are not lost. Future work is needed to disentangle these differences. Yet, our data showed that the cavin coat is flexible and can accommodate shapes from flat to highly curved.

Second, we find that EHD2 localizes to flat caveolae. EHD2 has been previously proposed to be associated with only the neck of caveolae^12,18^. Here, we observed both endogenous EHD2 or expressed EHD2-EGFP at flat caveolae. When comparing STED images of both caveolae types, EHD2 localization appears slightly more diffuse around flat caveolae (Fig. 5). Structural data of EHD2 and related EHD proteins (such as EHD4) demonstrated an ATP-dependent oligomerisation^47–50^ at lipid bilayers that forms a ring. Yet, it was recently shown that EHD can bind to flat membranes as a filament. These filaments change conformations to induce tubulation^49^. A more constricted EHD4 filament was observed when the underlying membrane curvature was increased. Therefore, we speculate that the accumulation of EHD2 around flat caveolae could be a nucleus or tether for its subsequent ring-like oligomerization at the caveolar neck. Together with pacsin2, which is also frequently detected at flat caveolae, membrane curvature could be generated. Notably, spherical caveolae showed a more confined EHD2 localization that suggests a denser ring-like structure near the caveolar neck. More detailed 3D atomic data is needed to fully understand how EHD2 oligomers (in concert with pacsin2) form at caveolae. Yet, these data again suggest that caveolae coats which include cavins and EHD2 are flexible polymers that can accommodate a range of curvatures and remain associated with each other and the membrane during these transitions.

We also investigated the localization of the relatively unstudied protein EHBP1 at caveolae. EHBP1 is known to bind to EHD2 proteins and actin^51–53^, and has been suggested to be involved in caveolae related processes^16,53^. EHBP1 was localized to only a subset of caveolae (Fig. 3). Morphological analysis revealed that it is more likely associated with curved caveolae. However, loss of EHBP1 did not alter caveolae number or shape (Fig. 6) indicating a regulatory rather than structural role of this protein.

From these data we conclude that flat and curved caveolae have similar core protein profiles (model Fig. 7I). This indicates that changes in membrane curvature at caveolae can occur without disassembly, re-assembly, or re-organization of the coat. Possibly, lipid changes drive caveolae shape changes as previously suggested^2,3,46,54,55^. We find that all major caveolae coat proteins (caveolins and cavins) can be found in both flat and invaginated caveolae to similar degrees. In line with this model, in the absence of EHD2, more vesicle-like caveolae were found. As EHD2 stabilizes caveolae at the plasma membrane, the loss of EHD2 results in highly mobile caveolae^11,12,17^ which would be reflected as an increase of spherical caveolae. Of note, similar to our observations, EHD2 deletion *in vivo* did not alter caveolae number in some organs^17,56^.

The proteins pacsin2 and EHBP1 were observed with more dispersed and sporadic localization profiles. Pacsin2 was mainly found at flat caveolae, where EHBP1 preferred bulb domains. This suggests that both proteins might be dynamically involved in the regulation of caveolae localization and traffic. Indeed, in the absence of pacsin2, caveolae appeared more flat (Fig. 6). In line with published results^13–15,20^ this indicates that pacsin2 is likely involved in caveolae curvature. Surprisingly, loss of pacsin2 and EHD2 combined (independently of EHBP1 levels) dramatically changed this result (Fig. 6) as the majority of caveolae became spheres. Furthermore, less caveolae were observed at the plasma membrane. Previous studies showed that caveolins and cavin proteins alone are able to form heterologous caveolae^57–60^ and, importantly, cellular uptake can occur^57^. Therefore, we speculate that the loss of EHD2, pacsin2, and EHBP1 leads to a unique caveolae structure. These “minimal” caveolae (containing caveolae coat proteins only) are much more mobile and less stable at the plasma membrane. This supports the idea that EHD2 restrains caveolae at the plasma membrane and pacsin2 is important for sphere formation.

Dynamin has been proposed to facilitate caveolae capture from the plasma membrane^19^. Surprisingly, across multiple experimental systems, we failed to localize dynamin to any caveolae shape (Fig. 7). Additionally, loss of all endogenous dynamins did not alter caveolae number or shape. Thus, a direct physical role for dynamin at caveolae is not supported by our data. In contrast, dynamin was clearly localized at nearby clathrin coated structures in abundance. How is caveolae fission achieved?

Besides its well-studied function in clathrin mediated endocytosis, dynamin can interact with actin^61–63^. Dynamin GTPase mutant K44A inhibits actin dynamics^63^. Caveolae are able to bind actin^64^ and when expressing the dynamin K44A mutant, caveolae mobility and endocytosis is inhibited^17,22,23,41^. Thus, we suggest that dynamin may be involved in actin-dependent caveolae traffic^64^. Possibly the combined functions of EHD2, pacsin2 and EHBP1 shapes and stabilizes the caveolar neck. Removal of these regulatory proteins shifts invaginated caveolae towards more highly curved spheres (as shown in triple KO/KD MEFs, Fig. 6). Binding of actin filaments to cavins or caveolins (via linker such as filamin A^64^) may then introduce additional mechanical force needed to overcome the energy barrier preventing membrane fission and endocytosis. Here, dynamin may form actin bundles with enhanced mechanical strength that allow the pulling of caveolae from the plasma membrane in a manner similar to membrane protrusions during cell fusion^61^ or clathrin-independent endocytosis^65^. Future studies are needed to clearly determine dynamin’s exact role in caveolae dynamics.

There are several specific limitations to our study. First, we focus on caveolae at the bottom (ventral) membrane of single cells. This is also true for most studies of caveolae that rely on evanescent field microscopy. Whether caveolae at the top membrane show similar protein composition and structural features will need to be determined in future studies. A similar issue exists for cells in complex tissues where they might be contacting other cells in three dimensions. Second, the resolution for STED of 40-60 nm is still too large to resolve any subtle sub-caveolae localization differences between the caveolae-associated proteins. If there is a slight heterogeneity of the eight proteins studied here, it will require much higher-resolution imaging methods such as cryoCLEM and cryo-tomography at the atomic scale. Third, we removed the unbound cytosol and nucleus with unroofing. While past studies have shown that this does not substantially change the plasma membrane or organelle structures in our systems, any subtle alterations to the underlying organelles will require higher-resolution and faster imaging studies in intact cells. Yet, as we see unexpected association (and not loss) of proteins at caveolae, we believe that the possible perturbations induced by unroofing do not impact our major experimental findings.

Here, we use correlative super-resolution STED and platinum replica correlative imaging to map the nanoscale location of caveolar proteins at caveolae of defined morphologies across entire plasma membranes of mammalian cells. By classifying proteins to both flat and curved caveolae, we generate a structural model for the molecular control of caveolae shape changes. We find that all the major caveolar coat proteins and the key ATPase EHD2 associates with both flat and curved caveolae to similar degrees. We propose that caveolae can dynamically flatten and curve as a single structural unit without loss or recruitment of the cavin coat or EHD2 (see model Fig. 7I). These are changes to the classical model of caveolae coat dynamics. Two other regulator proteins EHBP1 and pacsin2 differentially and sporadically associate with curved or flat sites. This suggest that these proteins act as regulators and not major architectural components. Knock-down/knockout of these regulators results in changes in the distribution of caveolae number and curvature, again, suggest a regulatory role for these proteins. Dynamin does not associate with caveolae. We propose a model where caveolae act as assembled malleable membrane curvature units that can capture excess membrane and extracellular lipids to regulate membrane tension or drive lipid transport in mammalian cells.

## Methods

### Cell culture

Wildtype, EHD2 knockout mouse embryonic fibroblasts (MEFs, previously described^17^) and Dynamin knockout MEFs (generously shared by Pietro De Camilli,^44,66^) were cultured in Dulbecco’s modified Eagle’s medium (DMEM, Gibco) supplemented with 10% fetal bovine serum (FBS) and 1% penicillin and streptomycin (Gibco). HUVEC (obtained from Promocell) were cultured in endothelial cell growth basal medium including SupplementMix (Promocell). Medium was changed every 2 days. For fluorescence or electron microscopy experiments cells were seeded on fibronectin coated glass dishes (#1.5 high precision, 25 mm; for STED: etched grid coverslip Bellco Biotechnology) and cultivated for 24 – 48 h at 37C in 5% CO_2_. MEFs were used for experiments until passage 35, HUVEC were used until passage 7. Triple Dynamin (Dyn1/2/3) knockout was induced by 1 μM 4-hydroxytamoxifen (Sigma) as previously described^44^. Briefly, MEFs were seeded sub-confluent and 1 μM 4-hydroxytamoxifen for 2 days was applied, followed by fresh DMEM containing 300 nM 4-hydroxytamoxifen for 4 days. Dynamin protein level was evaluated after 6 days by Western Blot and experiments were performed.

### Hypo-osmotic shock in HUVEC

Hypo-osmotic shock was induced by incubation of HUVEC in pre-warmed 1:5 (v%) growth medium diluted with water for 5 min. Afterwards HUVEC were immediately unroofed and fixed to prepare plasma membrane sheets. Evaluation of hypo-osmotic shock was done by phase contrast imaging inspecting cell shape over the time-course of 1 – 10 min.

### Plasmid transfection and siRNA treatment

Lipofectamine3000 (Invitrogen) was used to transfect MEFs seeded in 6 well plates (100.000 cells/well) with 2.5 μg plasmid accordingly to the manufacturers protocol. siRNA treatment was performed with Lipofectamine RNAiMax (Invitrogen), whereby the final siRNA concentration was 50 pmol per well (6 well plate). SMARTpool (mix of 4 siRNAs/target, Dharmacon) against mouse pacsin2 (#M-045093-01-0005) and mouse EHBP1 (#M-052068-01-0005) were used to obtain sufficient knockdown which was evaluated by Western blotting. All experiments were carried out after 48 h incubation. The following plasmids were used: pCaveolin1-EGFP, pCaveolin2-EGFP, pCavin1-EGFP, pCavin2-EGFP, pCavin3-EGFP, pPacsin2-EGFP, pEHD2-EGFP, pEHBP1-EGFP, pHis-Cavin1-EGFP, pDynamin2-GFP, pDynamin2-K44A-GFP.

### Preparation of plasma membrane sheets

Cells were unroofed prior to immunofluorescence staining or TEM preparation to obtain plasma membrane sheets as described previously (^37,67^). Briefly, cells seeded on glass dishes were washed with PBS and cell membrane stabilization buffer (70 mM KCl, 30 mM HEPES maintained at pH 7.4 with KOH, 5 mM MgCl_2_, 3 mM EGTA), and placed in fresh stabilization buffer. The unroofing was performed with 4% PFA (EM grade, freshly prepared, Electron Microscopy Science) that was splattered with a 19-gauge needle and syringe on the cells. Afterwards, the unroofed cells were placed in fresh 4% PFA (for immunofluorescence) or in 2% glutaraldehyde (for TEM, EM grade, Electron Microscopy Science) for fixation at 4C.

### Immunofluorescence staining and dyes

The unroofed cells were incubated in 3% bovine serum albumin/PBS (BSA, m/v, fresh) for 1.5 h, followed by primary antibody (1:100 in 3%BSA/PBS) incubation for 1 h. Next, cells were washed thoroughly with PBS and the secondary antibody tagged with fluorescence dye (1:500) or GFP-nanobody (1:500) was applied for 1 h. Afterwards, cells were washed 4 times in PBS and stored in fresh PBS at 4C until the samples were imaged. The following antibodies were used: anti-Caveolin1-Rabbit (abcam), anti-Caveolin1-mouse (SantaCruz), anti-Cavin1-Rabbit (abcam), anti-EHD2-goat (abcam), anti-Pacsin2-Rabbit (Proteintech), anti-EHBP1-Rabbit (Proteintech), anti-mouse-Clathrin light chain (Invitrogen, 1:2000), anti-mouse-Dynamin2 (Santa Cruz), anti-rabbit-Atto647N (Rockland), anti-mouse-Atto647N (Rockland), anti-goat-Atto647N (Rockland), Fab2-anti-rabbit-Alexa594 (ThermoFisher), Fab2-anti-mouse-Alexa488 (ThermoFisher), Fab2-anti-mouse-Alexa568 (ThermoFisher), GFP-booster-Atto647N (Chromotek), Phalloidin-Alexa488 (ThermoFisher).

### STED microscopy

Leica TCS SP8 microscope was used for 3 color gated STED with 100x objective (NA), including tunable white laser 470-670 nm, 775 and 592 nm depletion laser, and PMT and HyD Sp GaAsP detectors. The stained (unroofed) cells were imaged in PBS at room temperature. Depletion laser levels for Atto647N was between 25-50%, for Alexa594 between 40-75%, whereby Caveolin1 spot diameter size was used for STED evaluation. STED image size was 19.394 μm with a pixel size of 18.94 nm. Final lateral resolution was between 40 – 60 nm as determined with 40 nm fluorescent beads (Abberior).

### Gold labeling of cavin1 and caveolin1

Membrane sheets were prepared as described above, followed by fixation with 4% PFA for 20 min. Afterwards, the membranes were washed extensively with PBS (4-5x times), followed by two 0.1% EDTA/PBS (Sigma) washing steps and incubation with 3% BSA/PBS (m/v, fresh) for 1h. 10 nm Ni-NTA-Nanogold (Nanoprobes #2084) solution was diluted 1:5 in PBS and added to His-Cavin1-EGFP overexpressing MEF membrane sheets. The samples were first incubated for 15 min on orbital shaker followed by 45 min incubation without shaking. Next, the cells were treated similarly to Platinum replica as described below^68^. Caveolin1 was tagged with specific antibody (anti-Caveolin1-Rabbit, abcam, 1:100 in 3% BSA/PBS), and a secondary Rabbit antibody labelled with 12 nm gold particles (Dianova, 1:30 in 3% BSA/PBS) was applied. Investigation of gold labeling on Pt replica membrane sheets was done by TEM.

### Platinum replica preparation

The plasma membrane sheets were prepared for TEM as described previously^37^. Briefly, the unroofed cells were fixed in 2% glutaraldehyde for at least 20 min, followed by extensive washing with PBS and tannic acid (1 mg/ml dest. H_2_O) treatment for 20 min. Next, the cells were stained with 0.1% (v/v) uranyl acetate for 20 min. Afterwards, the membrane sheets were dehydrated by an increasing EtOH row (15% - 100%), followed by critical point drying with CO_2_. Platinum and carbon coating was carried out in Leica ACE900 freeze fracture in which, at first, 3 nm Pt and secondly 5.5 nm carbon was applied on the membrane sheets. The glass dish of the coated samples were removed by 10% (v/v) hydrofluoric acid, and the replicas were placed on Formvar/carbon coated 75 mesh EM copper grids (Ted Pella).

### TEM

TEM imaging was performed using a JOEL 1400 microscope at 15000 magnification (Pixel size 1.23 nm). Electron tomograms were obtained at 12000 magnification (Pixel size 1.56 nm) from - 60 to 60 degrees, with 1 degree increment. To obtain montage TEM images and tomograms SerialEM software was used^69^. Etomo/3DMOD was used to align tomogram stacks in fiducial-less mode with patch tracking, and IMOD was used for analysis^70^.

### STED-CLEM

Correlation of STED and TEM images was achieved by using gridded glass coverslips for correct cell assignment. After STED imaging a confocal tile scan of the grid was acquired including the cells of interest, followed by replica EM processing as described above. The region of interest was then cut out from the glass grid and phase contrast imaging (Nikon) of Pt replicas was used to re-assign the imaged cells on the TEM grids. After TEM imaging the STED images were aligned to the TEM montages by using a Matlab code previously described (^38,67^). Brief alignment was obtained by arranging the cell borders visible in both images. Exact alignment of STED and TEM images was achieved by clathrin fluorescence staining position to their distinctive clathrin structures in the TEM image, as well as Caveolin1 staining and caveolae structures in the replicas. Inspection and analysis of CLEM images was done in ImageJ/Fiji and Matlab accordingly to Sochacki et al. (2018).

### STORM-CLEM

STORM-CLEM data were all previously published in Sochacki et al., 2007^38^ with the exception of Dyn2-K44A expression in SK-MEL-2 cells. These data were acquired in the same manner as the previously published data using a Dyn2 (K44A)-GFP plasmid and staining with Alexa Fluor 647-conjugated GFP nanotrap^71^. The K44A point mutation was obtained with the Quickchange mutagenesis kit (Agilent) and the following primer set: FWD-TGG GCG GCC AGA GCG CCG GCG CGA GTT CGG TGC TCG AGA; REV-TCT CGA GCA CCG AAC TCG CGC CGG CGC TCT GGC CGC CCA. Sequence was confirmed following mutation.

### Protein isolation and Western Blotting

For protein isolation and Western blotting 100.000 – 200.000 cells were plated in 6 well plate and incubated for 48 h. After cells were washed with ice-cold PBS, 100 μl ice-cold RIPA buffer supplemented with proteases inhibitors (abcam) was added for cell lysis, and lysed cells were transferred in 1.5 ml tubes for vortexing. Next, cell lysates were incubated for 30 min on ice followed by 10 min centrifugation (13000 rpm) at 4C. 10 μl of the supernatant was used for SDS-PAGE on 8-12% Tris-Glycine gels (NOVEX™, Invitrogen) with Tris-Glycine SDS Running buffer (NOVEX™) at 120 V. Western Blotting was performed by MiniBlot (Biorad) with ready-to-use membranes (NOVEX™, Invitrogen) accordingly to the manufacturers protocol. Afterwards, membranes were incubated for 1 h at room temperature in 5% milk/TBS-T (1% Tween-20 diluted in TBS, NOVEX™). Primary antibodies were diluted in 5% milk/TBS-T and applied on the membranes over night at 4C (on horizontal shaker). After 3 times 10 min washing with TBS-T secondary antibody solution was added for 2h at room temperature. ECL solution (Amersham) was used for detection of protein levels in ChemiDoc XRS system (Biorad).

#### Antibodies

anti-Dynamin 1/2/3-mouse (BD Science, 1:1000), anti-Pacsin2-Rabbit (Proteintech, 1:500), anti-EHBP1-Rabbit (Proteintech, 1:1000), anti-GAPDH-Rabbit (Cell Signaling, 1:1000), anti-mouse-HRP (dianova, 1:5000), anti-Rabbit-HRP (dianova, 1:5000).

### Statistical analysis

All statistical analysis was carried out in Origin 2018b. First, data sets were analyzed by descriptive statistics (incl. mean, standard error of the mean (SE), median, min, max, standard derivation (SD), 5-95% interval) and normal distribution was tested by Shapiro-Wilk and Kolmogorov-Smirnov test. If data sets were normally distributed statistical differences were evaluated by two-tailed t-test, otherwise Mann Whitney test was applied (significance level 0.05, exact P value was measured). The following range of statistical differences is used in all figures: * P < 0.05, ** P < 0.01, *** P < 0.001, **** P < 0.0001.

## Acknowledgements

We thank the NHLBI electron microscopy and advanced light microscopy cores, and the FMP light and electron facility for their support and technical assistance. We thank Dr. Pietro De Camilli for the kind gift of the dynamin triple knockout MEFs. We thank Fabian Lukas and Tania López-Hernández for providing C2C12 myoblast cells and mouse astrocytes. We thank the Taraska lab for critical evaluation of experiments and manuscript.

## Author Contributions

C.M., M.L. and J.W.T. designed and discussed the experiments. C.M. performed and analyzed all experiments. K.A.S. wrote Matlab code for analysis of CLEM and STED data, supported and discussed analysis, performed STORM-CLEM and PREM of HeLa. A.D. did STORM-CLEM of Sk-mel2 cells. D.P. supported EM. V.H. supported and discussed experiments. C.M. and J.W.T. wrote the manuscript with input from all authors.

## Supplemental Figures and Videos

**Suppl. Figure S1:**
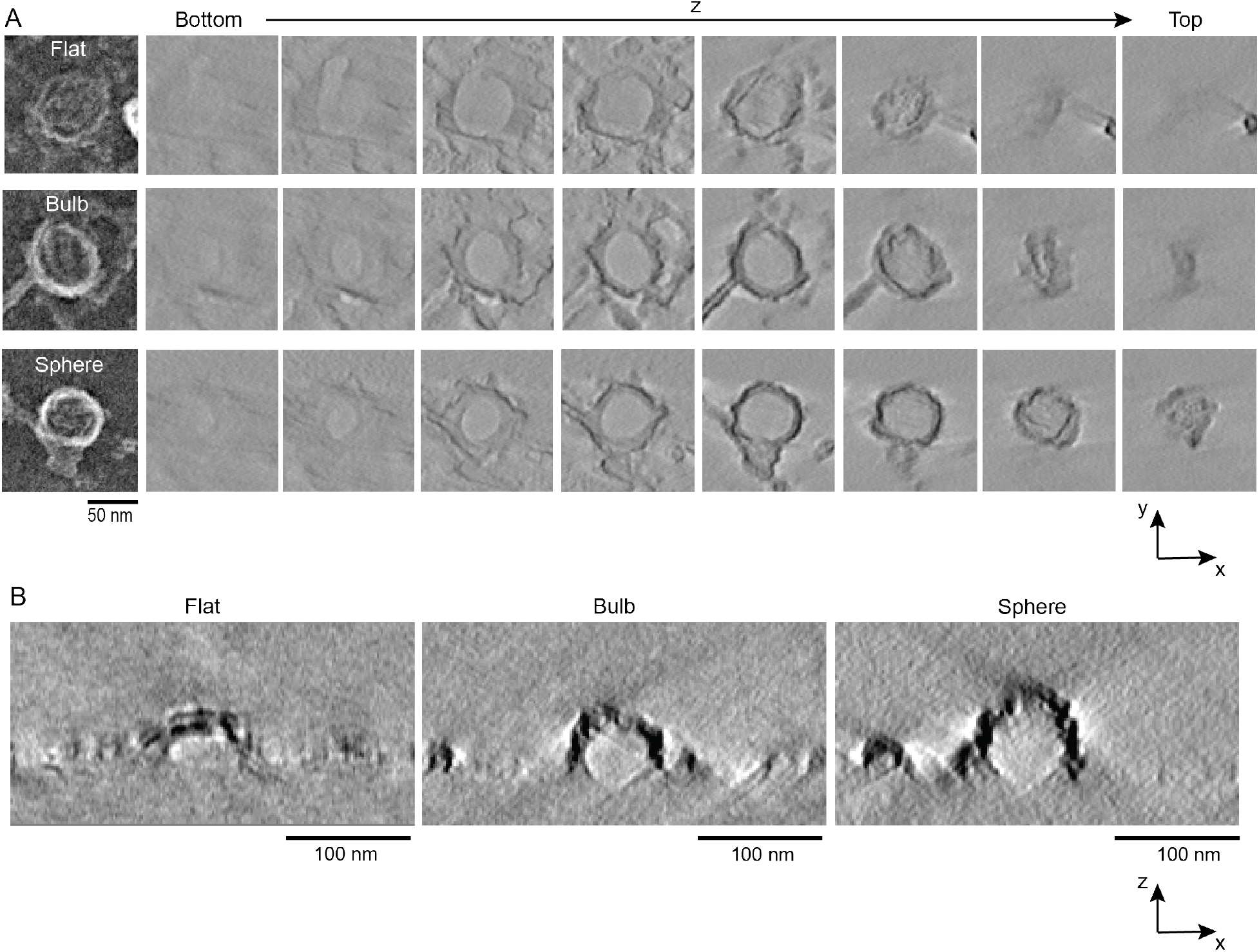
Distinct z-profiles for flat, bulb and spherical caveolae. (A) Representative TEM tomogram xy-images of caveolae subtypes (flat, bulb and sphere) from bottom to top. The same z slice is shown for each caveolae type. See also Suppl. Video S1-4. (B) TEM tomogram xz-images for flat, bulb, and spherical caveolae depicted in A.

**Suppl. Video1: Electron tomogram of MEF plasma membrane sheet including caveolae**.

**Suppl. Video2: Electron tomogram of flat caveolae (related to S1)**.

**Suppl. Video3: Electron tomogram of bulb-shaped caveolae (related to S1)**.

**Suppl. Video4: Electron tomogram of spherical caveolae (related to S1)**.

**Suppl. Figure S2:**
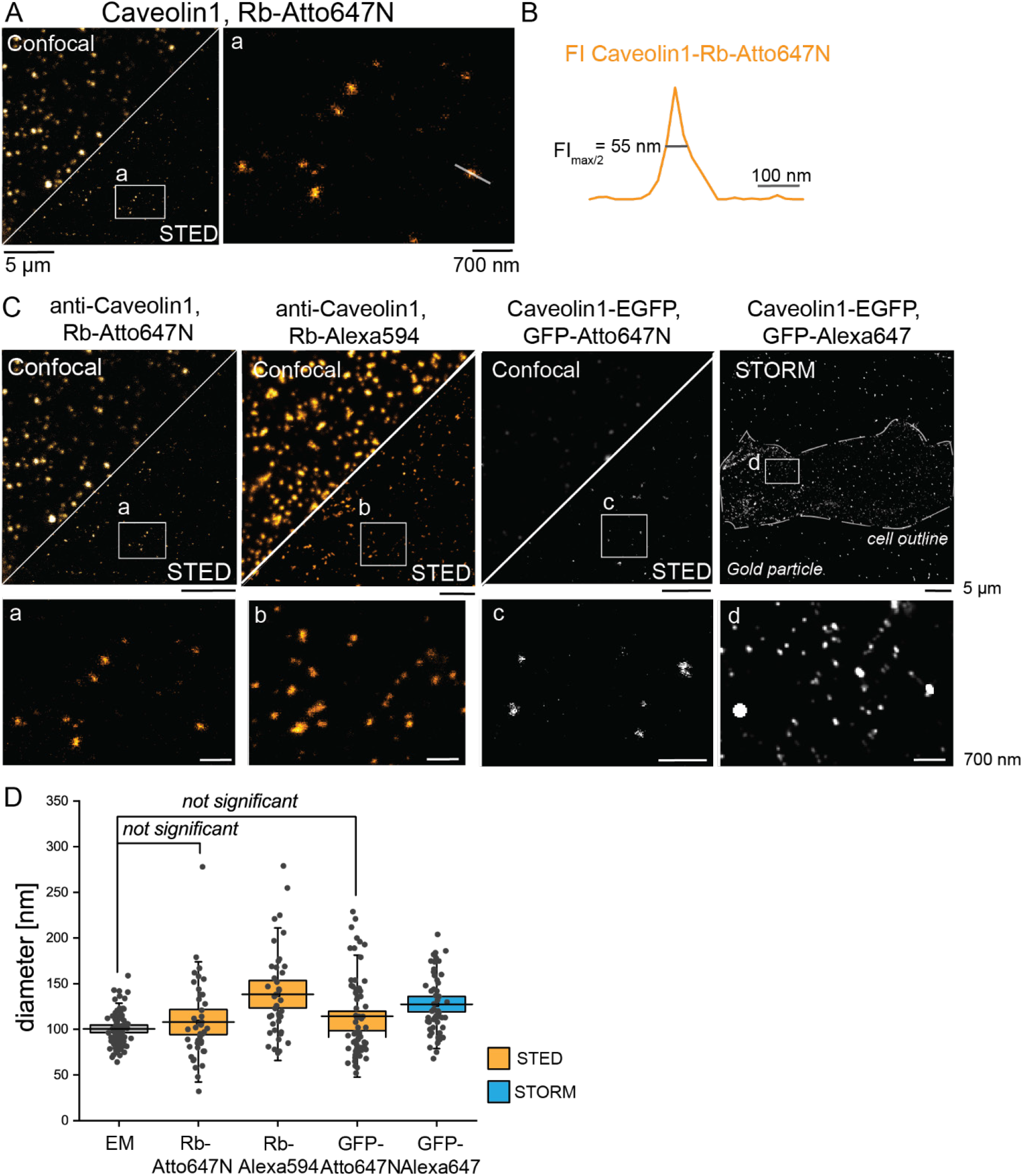
Exact caveolae size measurement by STED microscopy. (A) Representative STED image of endogenous caveolin1 immunolabelled and tagged by Atto647N (Rabbit-Atto647N) in MEF plasma membrane sheets. Enlarged selection (a) illustrates single caveolin1 spots. White line indicates plot profile shown in (B). (B) STED fluorescence plot profile (in orange) of a caveolin1 spot. FI_max/2_ = half maximum of fluorescence intensity spot. (C) Representative caveolin1 fluorescence images obtained by STED and STORM (Stochastic Optical Reconstruction Microscopy). Lower panel illustrates zoom selection of the individual STED or STORM images. (D) Caveolin1 spot size measurement based on fluorescence plot profiles. STED (orange) and STORM (blue) data was compared to diameter measured in PREM images (Fig. 1B). Box plot indicates mean ± SE, whiskers illustrate SD, each replicate is depicted (n(EM) = 82, n(STED-Rb-Atto647N) = 41, n(STED-Rb-Alexa594) = 42, n(STED-GFP nanobody-Atto647N) = 65, n(STORM-Alexa647) = 59). No statistical difference between EM and Atto647N (secondary antibody and GFP nanobody) was calculated by Mann Whitney test.

**Suppl. Figure S3:**
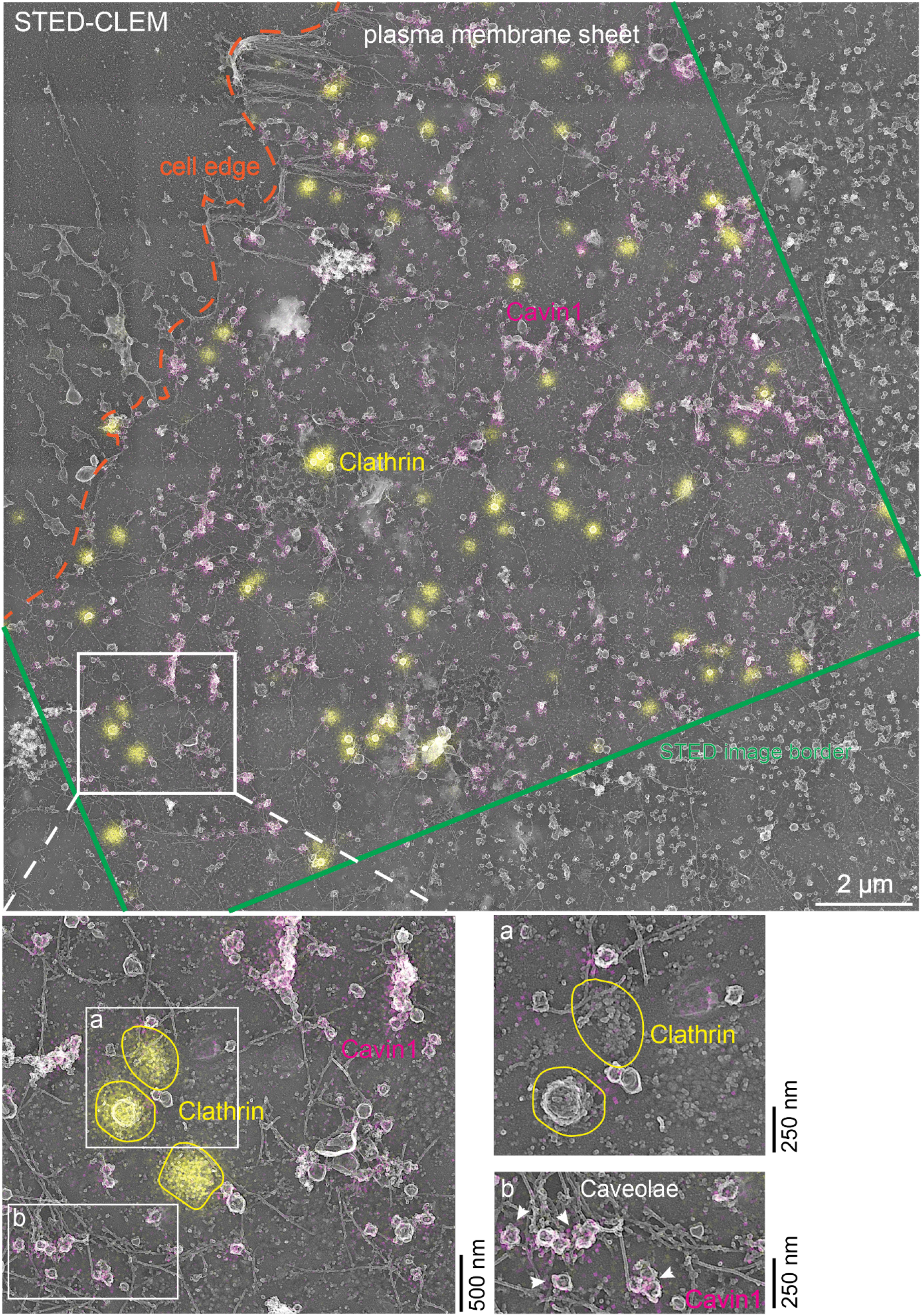
STED-CLEM overview of plasma membrane sheets showing clathrin and cavin1. Representative STED-CLEM image showing the acquired STED image that was correlated to the platinum replica TEM image. STED image border is shown in green, cell edge in orange. To achieve alignment of STED and PREM images, at first, cell edges were roughly aligned. Afterwards, detailed correlation was done by aligning clathrin clathrin fluorescence spots (in yellow) to the visible clathrin structures in the PREM image (done in Matlab, accordingly to^38,67^). Notably, due to the mass of the clathrin antibody, the characteristic geometric clathrin is partially obscured (see enlarged selection in a). Cavin1 fluorescence (in magenta) correlates to caveolae in PREM image (b, white arrows point to caveolae).

**Suppl. Figure S4:**
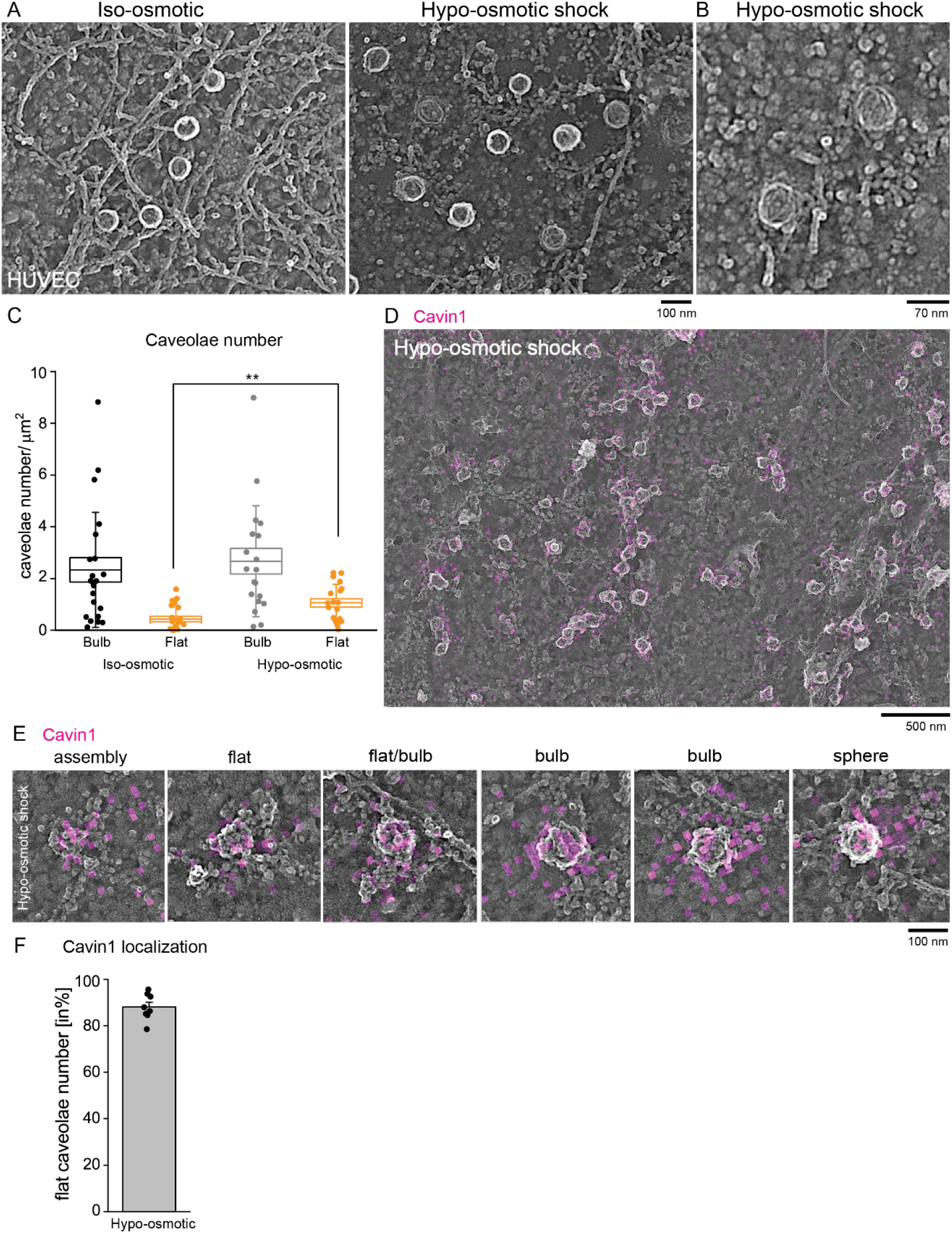
Cavin1 localization at flat and bulb caveolae after osmotic shock in HUVEC. (A-B) Representative PREM images of HUVEC plasma membrane in iso-osmotic control or after hypo-osmotic shock. (A) Number of invaginated (bulb and sphere) or flat caveolae at the plasma membrane in iso-osmotic HUVECs or after hypo-osmotic shock (n(iso-osmotic) = 22, n(hypo-osmotic) = 19; box plot illustrates mean ± SE, whiskers show SD, each replicate is depicted, Mann Whitney test for statistical difference, ** P<0.01). (B) Representative STED-CLEM image of cavin1 (magenta) after hypo-osmotic shock. (C) STED-CLEM of cavin1 (magenta) in different caveolae types after hypo-osmotic shock. (D) Percentage of cavin1 positive flat caveolae in STED-CLEM images after hypo-osmotic shock (n= 372 flat caveolae/8 cells; bar plot indicates mean ± SE, each replicate is shown).

**Suppl. Figure S5:**
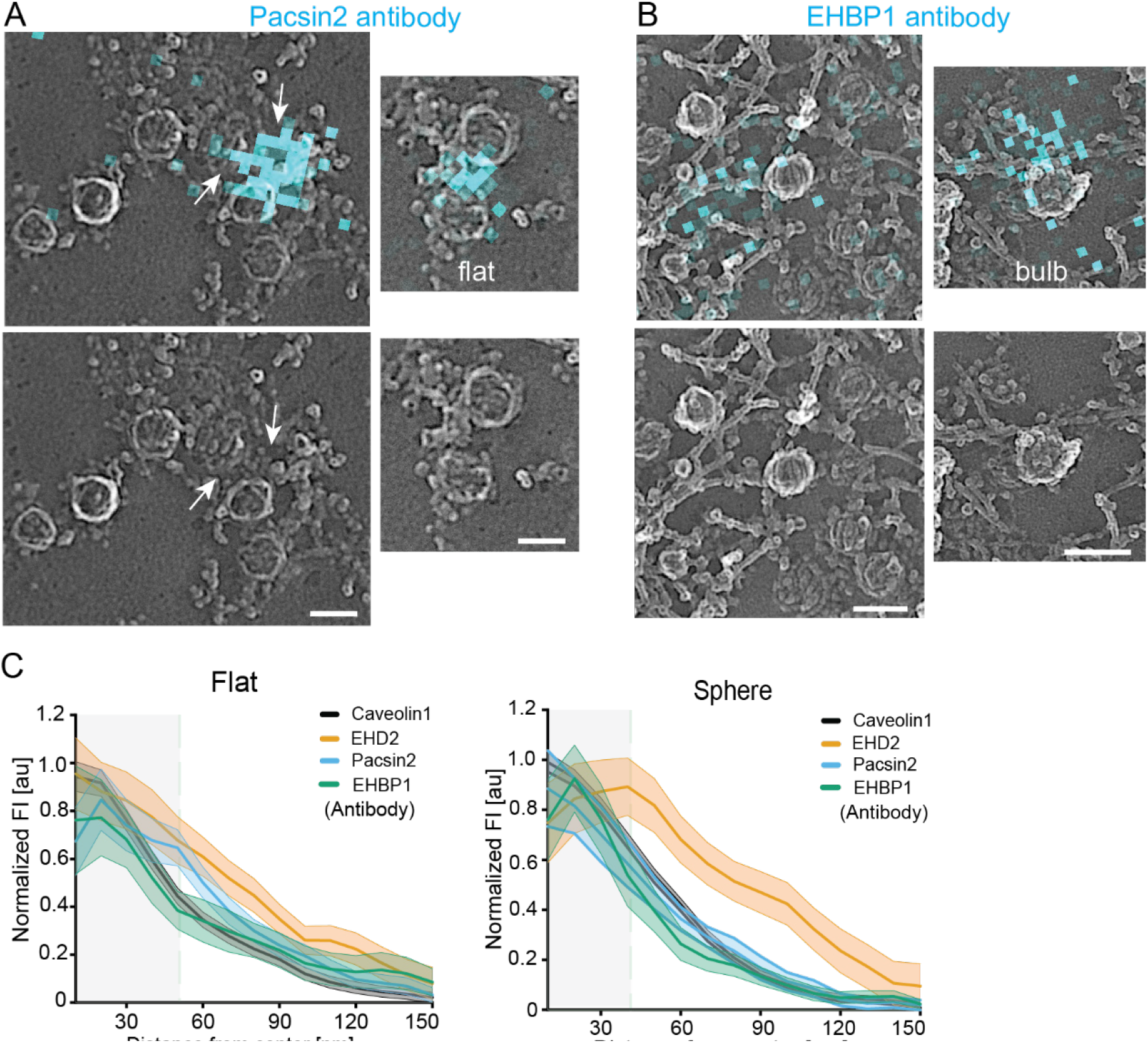
Endogenous staining of Pacsin2 and EHBP1 at caveolae. (A-B) Representative CLEM images for MEFs stained with antibodies against pacsin2 (A) or EHBP1 (B). The secondary antibody was labelled with Atto647N. White arrows in (A) indicate pacsin2 localization at flat caveolae. Scale bar shows 80 nm. (C) Quantitative analysis of flat or spherical caveolae by STED fluorescence profiles obtained from endogenous antibody staining. Line graphs show mean ± SE from the center of caveolae to their edges (indicated by green dashed line, caveolin1: n(flat) = 145, n(sphere) = 178; EHD2: n(flat) = 165, n(sphere) = 161; pacsin2: n(flat) = 77, n(sphere) = 49; EHBP1: n(flat) = 60, n(sphere) = 37).

**Suppl. Figure S6:**
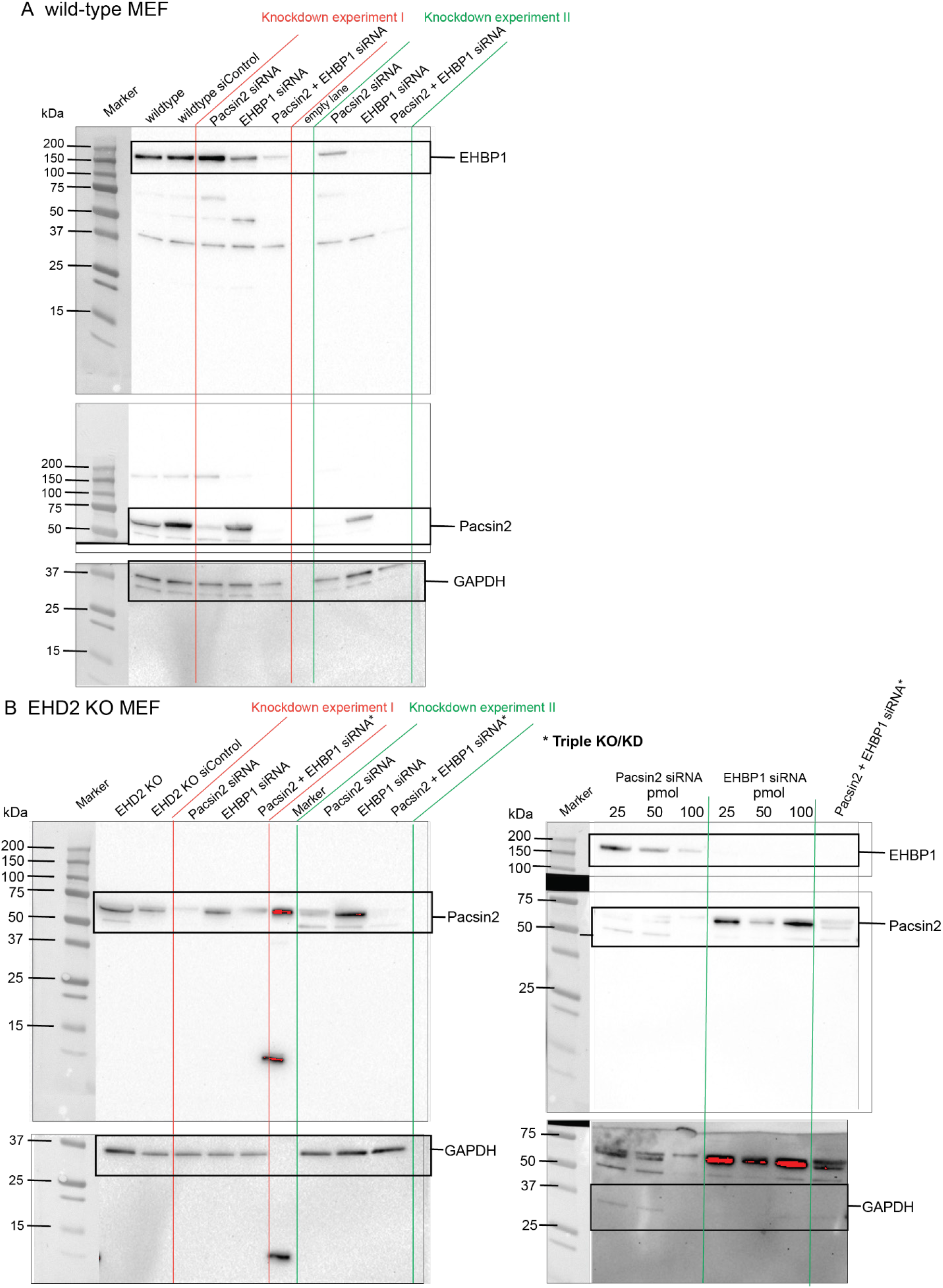
Western Blot validation of pacsin2 and EHBP1 knockdown. (A) Western Blot of pacsin2 and EHBP1 siRNA treated wild-type MEFs. Two independent siRNA knockdown experiments were blotted. EHBP1 antibody band size ∼ 160-180 kDa, pacsin2 antibody band size ∼ 60-65 kDa, GAPDH band size ∼37 kDa. (B) Western Blot of pacsin2 and EHBP1 siRNA treated EHD2 knockout (KO) MEFs. Two separate gels are shown.

**Suppl. Figure S7:**
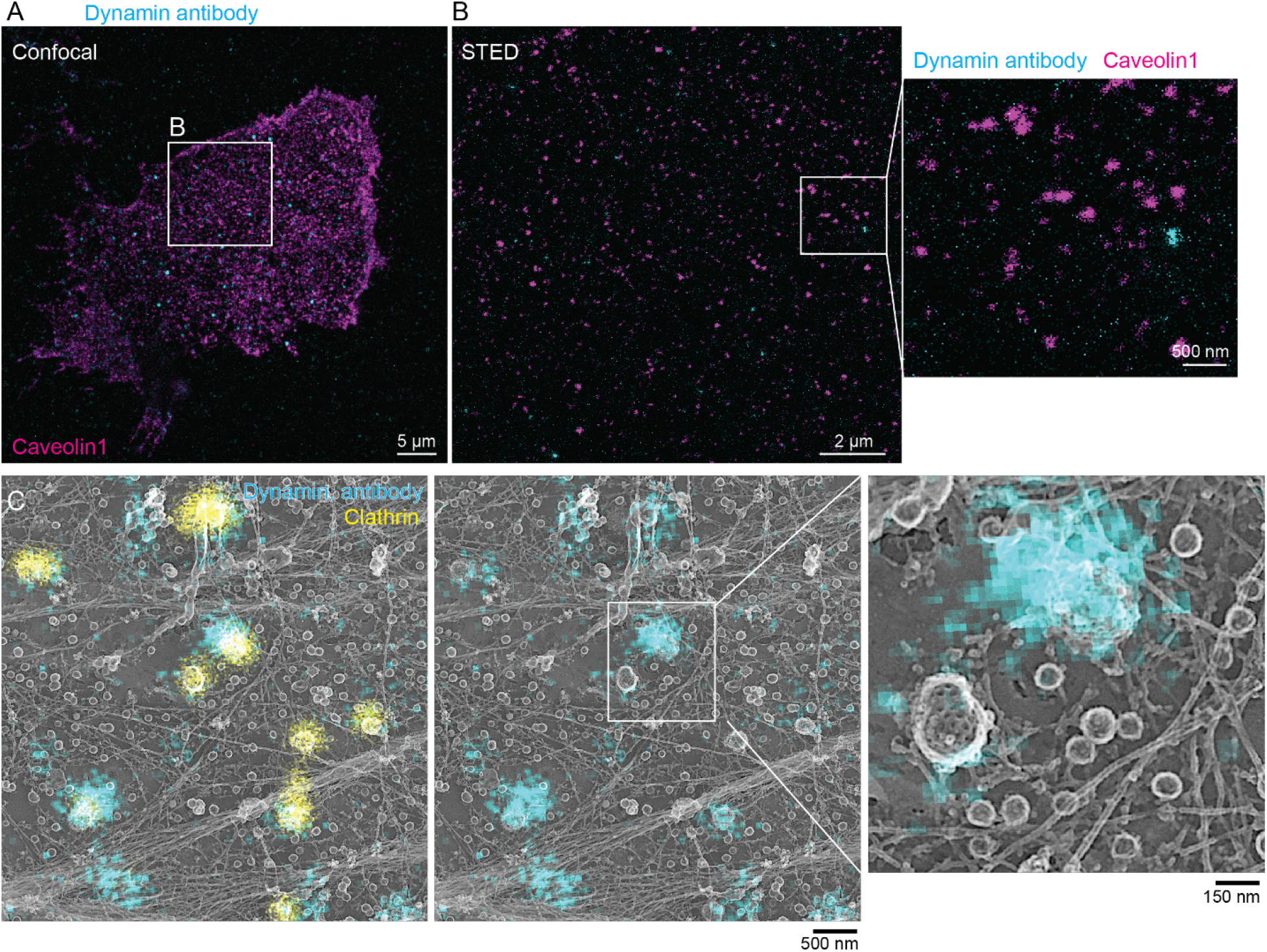
Immunostaining of endogenous dynamin in MEFs did not show localization to caveolae. (A) Confocal overview of MEF plasma membrane sheet stained against caveolin1 (magenta) and dynamin (cyan). Indicated selection was used for STED microscopy in B. (B) Representative STED image of caveolin1 (magenta) and dynamin (cyan) labelled with antibodies. (C) STED-PREM image showing antibody labelled dynamin (cyan) and clathrin (yellow). Enlarged panel illustrates caveolae and specific dynamin localization to clathrin coat.

**Suppl. Figure S8:**
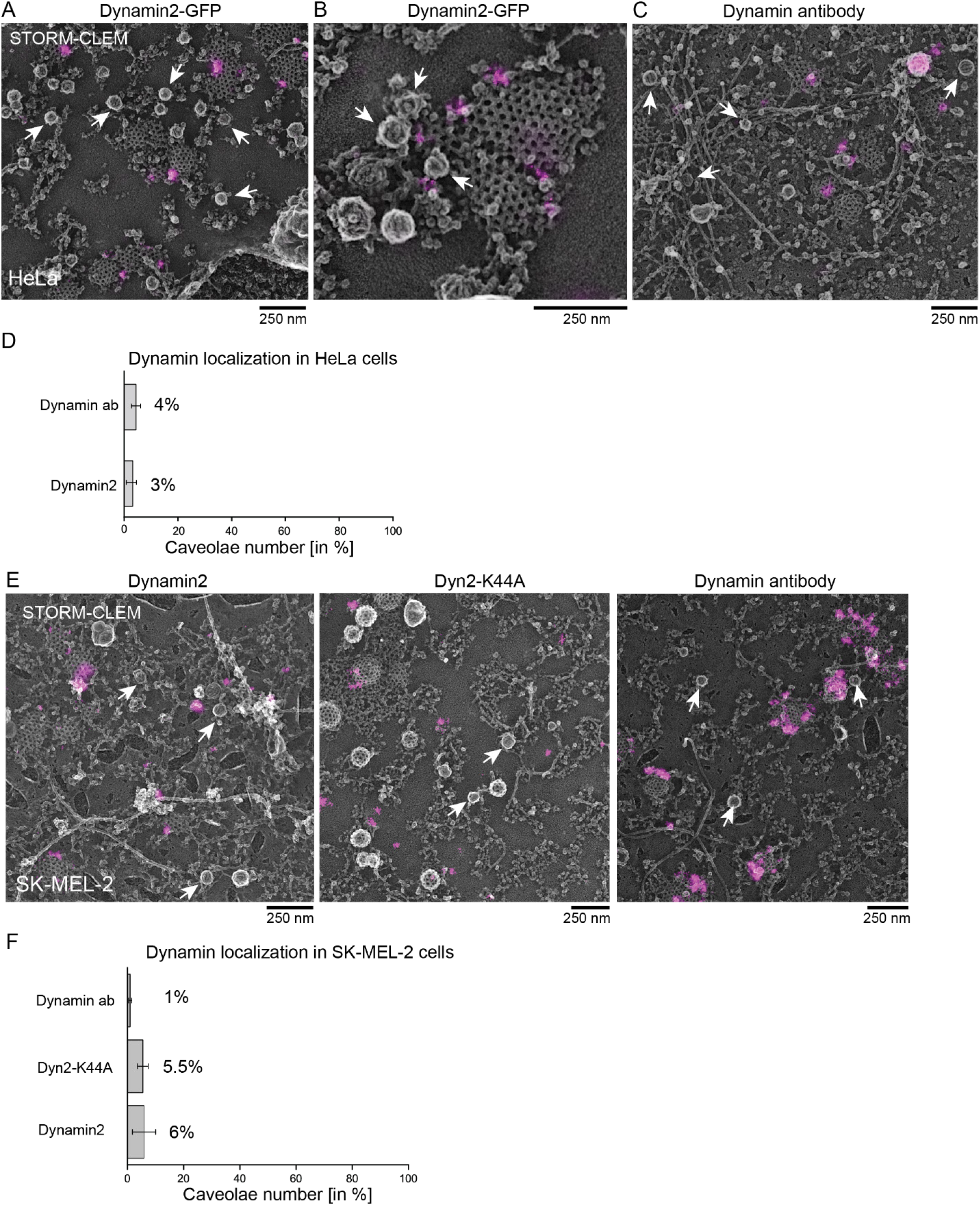
STORM-CLEM in HeLa and SK-MEL-2 cells did not show dynamin localization to caveolae. (A-B) Representative STORM-CLEM images illustrating STORM signal (in magenta) of dynamin2-GFP in Pt replicas of HeLa plasma membrane sheets. (B) illustrates dynamin localization to clathrin coat in close proximity to caveolae. White arrows indicate caveolae. (C) Representative STORM-CLEM images illustrating STORM signal (in magenta) of endogenous dynamin antibody labelling in Pt replicas of HeLa plasma membrane sheets. White arrows indicate caveolae. (D) Percentage of caveolae that were targeted by either dynamin2 or dynamin antibody in HeLa. Bar graph shows mean ± SE (n(Dynamin antibody) = 428/3 cells, n(Dynamin2) = 1649/7 cells). (E) Representative STORM-CLEM images illustrating STORM signal (in magenta) in SK-MEL-2 cells expressing either dynamin2-GFP or Dyn2-K44A-GFP, or were immune-stained to label endogenous dynamin. White arrows indicate caveolae. (F) Percentage of caveolae that were targeted by either dynamin2, dynamin2-K44A or dynamin antibody in SK-MEL-2 cells. Bar graph shows mean ± SE (n(Dynamin antibody) = 428/4 cells, n(Dyn2-K44A) = 270/4 cells, n(Dynamin2) = 426/5 cells).

**Suppl. Figure S9:**
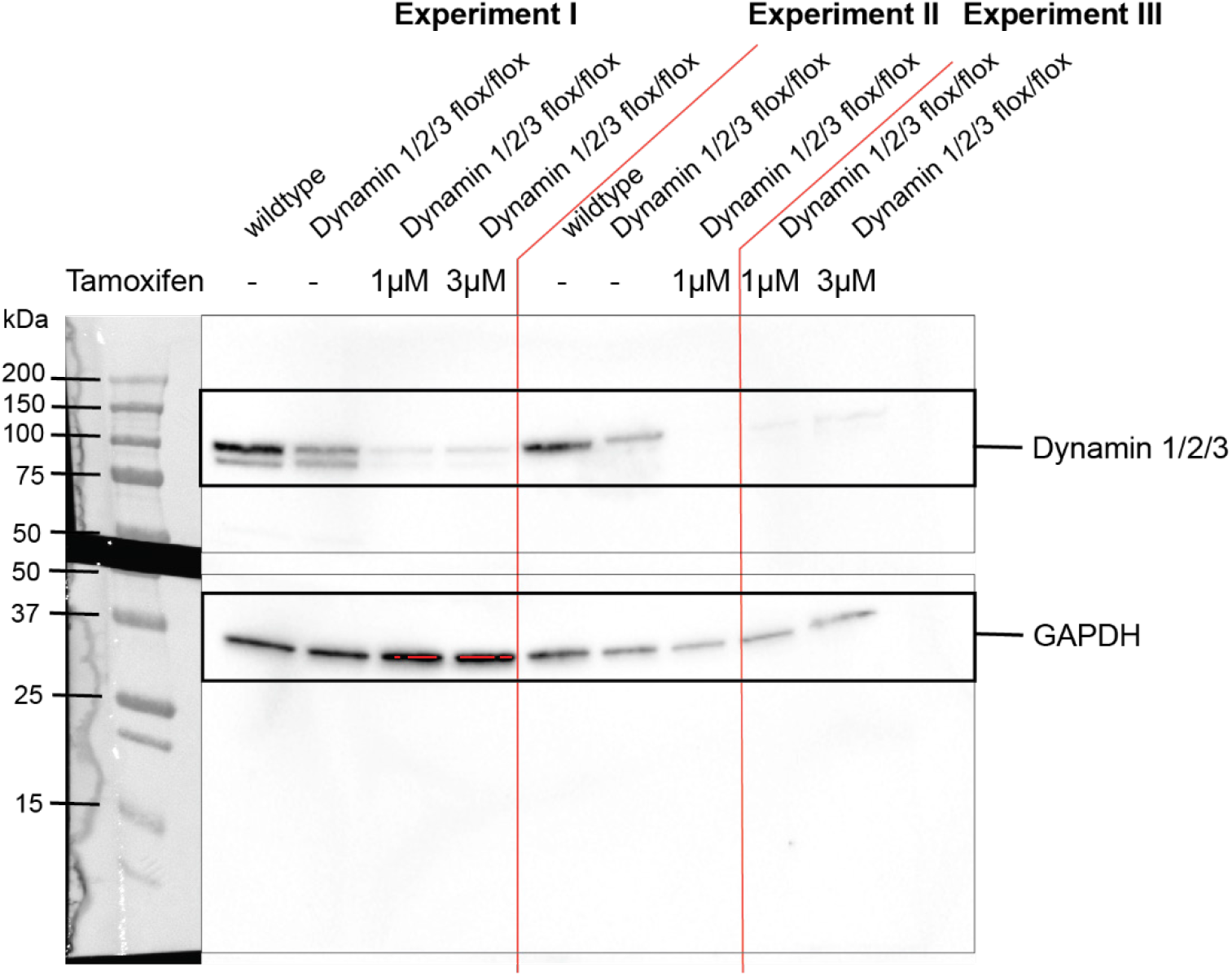
Western Blot validation of dynamin triple knockout. Western Blot illustrates three independent dynamin knockout experiments. Dynamin antibody recognizes all three isoforms, band size ∼ 100 kDa, GAPDH band size 37 kDa.

## References

1. Parton, R. G. & Simons, K. The multiple faces of caveolae. Nat. Rev. Mol. Cell Biol. 8, 185–94 (2007).

2. Zhou, Y. et al. Caveolin-1 and cavin1 act synergistically to generate a unique lipid environment in caveolae. J. Cell Biol. 220, (2021).

3. Parton, R. G., Kozlov, M. M. & Ariotti, N. Caveolae and lipid sorting: Shaping the cellular response to stress. J. Cell Biol. 219, 1–13 (2020).

4. Cheng, J. P. X. & Nichols, B. J. Caveolae : One Function or Many ? Trends Cell Biol. 26, 177–189 (2016).

5. Le Lay, S. & Kurzchalia, T. V. Getting rid of caveolins: Phenotypes of caveolin-deficient animals. Biochim. Biophys. Acta - Mol. Cell Res. 1746, 322–333 (2005).

6. Dewulf, M. et al. Dystrophy-associated caveolin-3 mutations reveal that caveolae couple IL6/STAT3 signaling with mechanosensing in human muscle cells. Nat. Commun. 10, 1–13 (2019).

7. Martinez-Outschoorn, U. E., Sotgia, F. & Lisanti, M. P. Caveolae and signalling in cancer. Nat. Rev. Cancer 15, 225–237 (2015).

8. Stoeber, M. et al. Model for the architecture of caveolae based on a flexible, net-like assembly of Cavin1 and Caveolin discs. Proc. Natl. Acad. Sci. U. S. A. 201616838 (2016) doi:10.1073/pnas.1616838113.

9. Kovtun, O., Tillu, V. A., Ariotti, N., Parton, R. G. & Collins, B. M. Cavin family proteins and the assembly of caveolae. J. Cell Sci. 128, 1269–1278 (2015).

10. Gambin, Y. et al. Single-molecule analysis reveals self assembly and nanoscale segregation of two distinct cavin subcomplexes on caveolae. Elife 3, 1–18 (2014).

11. Stoeber, M. et al. Oligomers of the ATPase EHD2 confine caveolae to the plasma membrane through association with actin. EMBO J. 31, 2350–2364 (2012).

12. Morén, B. et al. EHD2 regulates caveolar dynamics via ATP-driven targeting and oligomerization. Mol. Biol. Cell 23, 1316–29 (2012).

13. Senju, Y., Itoh, Y., Takano, K., Hamada, S. & Suetsugu, S. Essential role of PACSIN2/syndapin-II in caveolae membrane sculpting. J. Cell Sci. 124, 2032–2040 (2011).

14. Hansen, C. G., Howard, G. & Nichols, B. J. Pacsin 2 is recruited to caveolae and functions in caveolar biogenesis. J. Cell Sci. 124, 2777–2785 (2011).

15. Koch, D., Westermann, M., Kessels, M. M. & Qualmann, B. Ultrastructural freeze-fracture immunolabeling identifies plasma membrane-localized syndapin II as a crucial factor in shaping caveolae. Histochem. Cell Biol. 138, 215–230 (2012).

16. Webb, A. et al. EHBP1 and EHD2 regulate Dll4 caveolin-mediated endocytosis during blood vessel development. bioRxiv (2020) doi:10.1101/2020.05.19.104547.

17. Matthaeus, C. et al. EHD2-mediated restriction of caveolar dynamics regulates cellular fatty acid uptake. Proc. Natl. Acad. Sci. 117, 7471–7481 (2020).

18. Ludwig, A. et al. Molecular Composition and Ultrastructure of the Caveolar Coat Complex. PLoS Biol. 11, (2013).

19. Matthaeus, C. & Taraska, J. W. Energy and Dynamics of Caveolae Trafficking. Front. Cell Dev. Biol. 8, (2021).

20. Seemann, E. et al. Deciphering caveolar functions by syndapin III KO-mediated impairment of caveolar invagination. Elife 6, 1–37 (2017).

21. Senju, Y. et al. Phosphorylation of PACSIN2 by protein kinase C triggers the removal of caveolae from the plasma membrane. J. Cell Sci. 128, 2766–2780 (2015).

22. Oh, P., McIntosh, D. P. & Schnitzer, J. E. Dynamin at the neck of caveolae mediates their budding to form transport vesicles by GTP-driven fission from the plasma membrane of endothelium. J. Cell Biol. 141, 101–114 (1998).

23. Yao, Q. et al. Caveolin-1 interacts directly with dynamin-2. J. Mol. Biol. 348, 491–501 (2005).

24. Henley, J. R., Krueger, E. W. A., Oswald, B. J. & McNiven, M. A. Dynamin-mediated internalization of caveolae. J. Cell Biol. 141, 85–99 (1998).

25. Yamaguchi, T. et al. ROR1 sustains caveolae and survival signalling as a scaffold of cavin-1 and caveolin-1. Nat. Commun. 7, (2016).

26. Echarri, A. et al. An Abl-FBP17 mechanosensing system couples local plasma membrane curvature and stress fiber remodeling during mechanoadaptation. Nat. Commun. 10, (2019).

27. Golani, G., Ariotti, N., Parton, R. G. & Kozlov, M. M. Membrane Curvature and Tension Control the Formation and Collapse of Caveolar Superstructures. Dev. Cell 48, 523-538.e4 (2019).

28. Parton, R. G. et al. Caveolae: The FAQs. Traffic 21, 181–185 (2020).

29. Parton, R. G., Tillu, V., McMahon, K. A. & Collins, B. M. Key phases in the formation of caveolae. Curr. Opin. Cell Biol. 71, 7–14 (2021).

30. Han, B. et al. Structure and assembly of CAV1 8S complexes revealed by single particle electron microscopy. Sci. Adv. 6, eabc6185.2 (2020).

31. Ariotti, N. et al. Molecular characterization of caveolin-induced membrane curvature. J. Biol. Chem. 290, 24875–24890 (2015).

32. Porta, J. C. et al. Molecular architecture of the human caveolin-1 complex. bioRxiv (2022) doi:10.1101/2022.02.17.480763.

33. Sinha, B. et al. Cells respond to mechanical stress by rapid disassembly of caveolae. Cell 144, 402–413 (2011).

34. Parton, R. G., McMahon, K. A. & Wu, Y. Caveolae: Formation, dynamics, and function. Curr. Opin. Cell Biol. 65, 8–16 (2020).

35. Torrino, S. et al. EHD2 is a mechanotransducer connecting caveolae dynamics with gene transcription. J. Cell Biol. 1–14 (2018).

36. Del Pozo, M. A., Lolo, F.-N. & Echarri, A. Caveolae: Mechanosensing and mechanotransduction devices linking membrane trafficking to mechanoadaptation. Curr. Opin. Cell Biol. 68, 113–123 (2021).

37. Sochacki, K. A. & Taraska, J. W. Correlative fluorescent super-resolution localization microscopy and platinum replica EM on unroofed cells. Methods Mol. Biol. 1663, 231–252 (2017).

38. Sochacki, K. A., Dickey, A. M., Strub, M. P. & Taraska, J. W. Endocytic proteins are partitioned at the edge of the clathrin lattice in mammalian cells. Nat. Cell Biol. 19, 352–361 (2017).

39. Rothberg, K. G. et al. Caveolin, a Protein-Component of Caveolae Membrane Coats. Cell 68, 673– 682 (1992).

40. Sochacki, K. A. et al. The structure and spontaneous curvature of clathrin lattices at the plasma membrane. Dev. Cell 56, 1131-1146.e3 (2021).

41. Oh, P., Horner, T., Witkiewicz, H. & Schnitzer, J. E. Endothelin induces rapid, dynamin-mediated budding of endothelial caveolae rich in ET-B. J. Biol. Chem. 287, 17353–17362 (2012).

42. Ferguson, S. M. & De Camilli, P. Dynamin, a membrane-remodelling GTPase. Nat. Rev. Mol. Cell Biol. 13, 75–88 (2012).

43. Le, P. U., Guay, G., Altschuler, Y. & Nabi, I. R. Caveolin-1 is a negative regulator of caveolae-mediated endocytosis to the endoplasmic reticulum. J. Biol. Chem. 277, 3371–3379 (2002).

44. Park, R. J. et al. Dynamin triple knockout cells reveal off target effects of commonly used dynamin inhibitors. J. Cell Sci. 126, 5305–5312 (2013).

45. Shvets, E., Bitsikas, V., Howard, G., Hansen, C. G. & Nichols, B. J. Dynamic caveolae exclude bulk membrane proteins and are required for sorting of excess glycosphingolipids. Nat. Commun. 6, (2015).

46. Hubert, M. et al. Lipid accumulation controls the balance between surface connection and scission of caveolae. Elife 9:e55038 (2020) doi:10.7554/eLife.55038.

47. Daumke, O. et al. Architectural and mechanistic insights into an EHD ATPase involved in membrane remodelling. Nature 449, 923–7 (2007).

48. Melo, A. A. et al. Structural insights into the activation mechanism of dynamin-like EHD ATPases. Proc. Natl. Acad. Sci. 114, 5629–5634 (2017).

49. Melo, A. A. et al. Cryo-electron tomography reveals structural insights into the membrane binding and remodeling activity of dynamin-like EHDs. bioRxiv 2021.09.20.461157 (2021).

50. Hoernke, M. et al. EHD2 restrains dynamics of caveolae by an ATP-dependent, membrane-bound, open conformation. Proc. Natl. Acad. Sci. U. S. A. 114, E4360–E4369 (2017).

51. Guilherme, A., Soriano, N. A., Furcinitti, P. S. & Czech, M. P. Role of EHD1 and EHBP1 in perinuclear sorting and insulin-regulated GLUT4 recycling in 3T3-L1 adipocytes. J. Biol. Chem. 279, 40062–40075 (2004).

52. Li, Z. et al. A novel Rab10-EHBP1-EHD2 complex essential for the autophagic engulfment of lipid droplets. Sci. Adv. 2, 1–16 (2016).

53. Simone, L. C., Caplan, S. & Naslavsky, N. Role of Phosphatidylinositol 4,5-Bisphosphate in Regulating EHD2 Plasma Membrane Localization. PLoS One 8, (2013).

54. Ariotti, N. et al. Caveolae regulate the nanoscale organization of the plasma membrane to remotely control Ras signaling. J. Cell Biol. 204, 777–792 (2014).

55. Fujita, A., Cheng, J., Tauchi-Sato, K., Takenawa, T. & Fujimoto, T. A distinct pool of phosphatidylinositol 4,5-bisphosphate in caveolae revealed by a nanoscale labeling technique. Proc. Natl. Acad. Sci. U. S. A. 106, 9256–9261 (2009).

56. Matthaeus, C. et al. ENOS-NO-induced small blood vessel relaxation requires EHD2-dependent caveolae stabilization. PLoS One 14, 1–22 (2019).

57. Walser, P. J. et al. Constitutive formation of caveolae in a bacterium. Cell 150, 752–763 (2012).

58. Jung, W. R. et al. Cell-free formation and interactome analysis of caveolae. J. Cell Biol. 217, 2141– 2165 (2018).

59. Tillu, V. A. et al. Cavin1 intrinsically disordered domains are essential for fuzzy electrostatic interactions and caveola formation. Nat. Commun. 12, 1–18 (2021).

60. Hill, M. M. et al. PTRF-Cavin, a Conserved Cytoplasmic Protein Required for Caveola Formation and Function. Cell 132, 113–124 (2008).

61. Zhang, R. et al. Dynamin regulates the dynamics and mechanical strength of the actin cytoskeleton as a multifilament actin-bundling protein. Nat. Cell Biol. 22, 674–688 (2020).

62. Lee, E. & De Camilli, P. Dynamin at actin tails. Proc. Natl. Acad. Sci. U. S. A. 99, 161–166 (2002).

63. Orth, J. D., Krueger, E. W., Cao, H. & McNiven, M. A. The large GTpase dynamin regulates actin comet formation and movement in living cells. Proc. Natl. Acad. Sci. U. S. A. 99, 167–172 (2002).

64. Echarri, A. & Del Pozo, M. A. Caveolae - mechanosensitive membrane invaginations linked to actin filaments. J. Cell Sci. 128, 2747–2758 (2015).

65. Johannes, L., Parton, R. G., Bassereau, P. & Mayor, S. Building endocytic pits without clathrin. Nat. Rev. Mol. Cell Biol. 16, 311–321 (2015).

66. Ferguson, S. et al. Coordinated Actions of Actin and BAR Proteins Upstream of Dynamin at Endocytic Clathrin-Coated Pits. Dev. Cell 17, 811–822 (2009).

67. Sochacki, K. A., Shtengel, G., Van Engelenburg, S. B., Hess, H. F. & Taraska, J. W. Correlative super-resolution fluorescence and metal-replica transmission electron microscopy. Nat. Methods 11, 305–308 (2014).

68. Prasai, B. et al. The nanoscale molecular morphology of docked exocytic dense-core vesicles in neuroendocrine cells. Nat. Commun. 12, 1–14 (2021).

69. Mastronarde, D. N. Automated electron microscope tomography using robust prediction of specimen movements. J. Struct. Biol. 152, 36–51 (2005).

70. Kremer, J. R., Mastronarde, D. N. & McIntosh, J. R. Computer visualization of three-dimensional image data using IMOD. J. Struct. Biol. 116, 71–76 (1996).

71. Rothbauer, U. et al. A versatile nanotrap for biochemical and functional studies with fluorescent fusion proteins. Mol. Cell. Proteomics 7, 282–289 (2008).

